# ADNP promotes neural differentiation by modulating Wnt/β-catenin signaling

**DOI:** 10.1101/761064

**Authors:** Xiaoyun Sun, Xixia Peng, Yuqing Cao, Yan Zhou, Yuhua Sun

## Abstract

ADNP (Activity Dependent Neuroprotective Protein) is proposed as a neuroprotective protein whose aberrant expression has been frequently linked to neural developmental disorders, including the Helsmoortel-Van der Aa syndrome. However, its role in neural development and pathology remains unclear. Using mESC (mouse embryonic stem cell) directional neural differentiation as a model, we show that ADNP is required for ESC neural induction and neuronal differentiation by maintaining Wnt signaling. Mechanistically, ADNP functions to maintain the proper protein levels of β-Catenin through binding to its armadillo domain which prevents its association with key components of the degradation complex: Axin and APC. Loss of ADNP promotes the formation of the degradation complex and hyperphosphorylation of β-Catenin by GSK3β and subsequent degradation via ubiquitin-proteasome pathway, resulting in down-regulation of key neuroectoderm developmental genes. We further show that ADNP plays key role in cerebellar neuron formation. Finally, *adnp* gene disruption in zebrafish embryos recapitulates key features of the mouse phenotype, including the reduced Wnt signaling, defective embryonic cerebral neuron formation and the massive neuron death. Thus, our work provides important insights into the role of ADNP in neural development and the pathology of the Helsmoortel-Van der Aa syndrome caused by *ADNP* gene mutation.

## Introduction

ADNP was first described as a neural protective protein and has been implicated in various neural developmental disorders and cancers^1^. This protein contains nine zinc fingers, a homeobox domain and a PxLxP motif, suggesting that it functions as a transcription factor. The exact roles of ADNP remain unclear, but an increasing number of studies have shown that it functions as a key chromatin regulator by interacting with chromatin remodelers or regulators. For instances, ADNP interacts with core sub-units of SWI/SNF chromatin remodeling complex such as BRG1 and BAF250^2^. By association with the chromatin regulator HP1, ADNP localizes to pericentromeric heterochromatin regions where it silences major satellite repeat elements^3^. Recently, ADNP was shown to form a triplex with CHD4 and HP1 to control the expression of lineage specifying genes in embryonic stem cells^4^.

Mouse *Adnp* mRNA is abundantly expressed during early gestation and reaches its maximum expression on E9.5. *Adnp-/-* mice are early embryonic lethal, displaying severe defects in neural tube closure and brain formation^5^. Of note, E8.5 *Adnp* mutant embryos exhibited ectopic expression of pluripotency gene *Pou5f1* and reduced expression of neural developmental gene *Pax6* in the anterior neural plate. These observations strongly suggested that ADNP plays essential roles during neurogenesis of mouse embryos.

De novo mutations in *ADNP* gene have recently been linked to neural developmental disorders, including the Helsmoortel-Van der Aa syndrome^6^. Patients display multiple symptoms, usually manifesting in early childhood with features such as intellectual disability, facial dysmorphism, motor dysfunction and developmental delay, which share many features with another neural developmental disorder: autism spectrum disorder (ASD). In fact, *ADNP* has been proposed as one of the most frequent ASD-associated genes as it is mutated in at least 0.17% of ASD cases^7^. In a mouse model, haploinsufficiency of *Adnp* leads to defective neuronal/glial formation and compromised cognitive function^8^. Due to the significant relevance of *ADNP* to neural developmental disorders, it is of great interests and importance in the field to elucidate the pathogenic mechanism by *ADNP* gene mutation^6^.

In this work, we hypothesized that loss of ADNP leads to neural developmental defects. We took use of ES cell directional neural differentiation as a model system to investigate the role of ADNP in embryonic neural development. We showed that ADNP is required for proper neural induction by modulating Wnt/β-catenin signaling. Mechanistically, ADNP functions to maintain protein levels of β-Catenin through binding to its armadillo domain which prevents its interaction with key components of degradation complex: Axin and APC. Loss of ADNP leads to hyperphosphorylation of β-Catenin by GSK3β and subsequent degradation via ubiquitin-proteasome pathway, resulting in down-regulation of neuroectoderm developmental genes. Small molecule-mediated activation of Wnt signaling can rescue the defects by loss of ADNP. This work provides important insights into the role of ADNP in neural development which would be useful for understanding the pathology of the Helsmoortel-Van der Aa syndrome caused by *ADNP* mutation.

## Results

### Generation and characterization of *Adnp-/-* ESCs

To understand the molecular function of ADNP, we generated *Adnp* mutant ESCs by using CRISPR/Cas9 technology. gRNAs were designed to target the 3’ end of exon 4 of mouse *Adnp* gene (Fig. 1a). We have successfully generated 2 *Adnp* mutant alleles (-4 and -5 bp), revealed by DNA genotyping around the CRISPR targeting site (Fig. 1b). ADNP protein was hardly detectable in *Adnp*-/- ESCs by Western blot using ADNP antibodies from different resources (Fig. 1c), which strongly indicated that the mutant alleles are functional nulls.

**Fig. 1.**
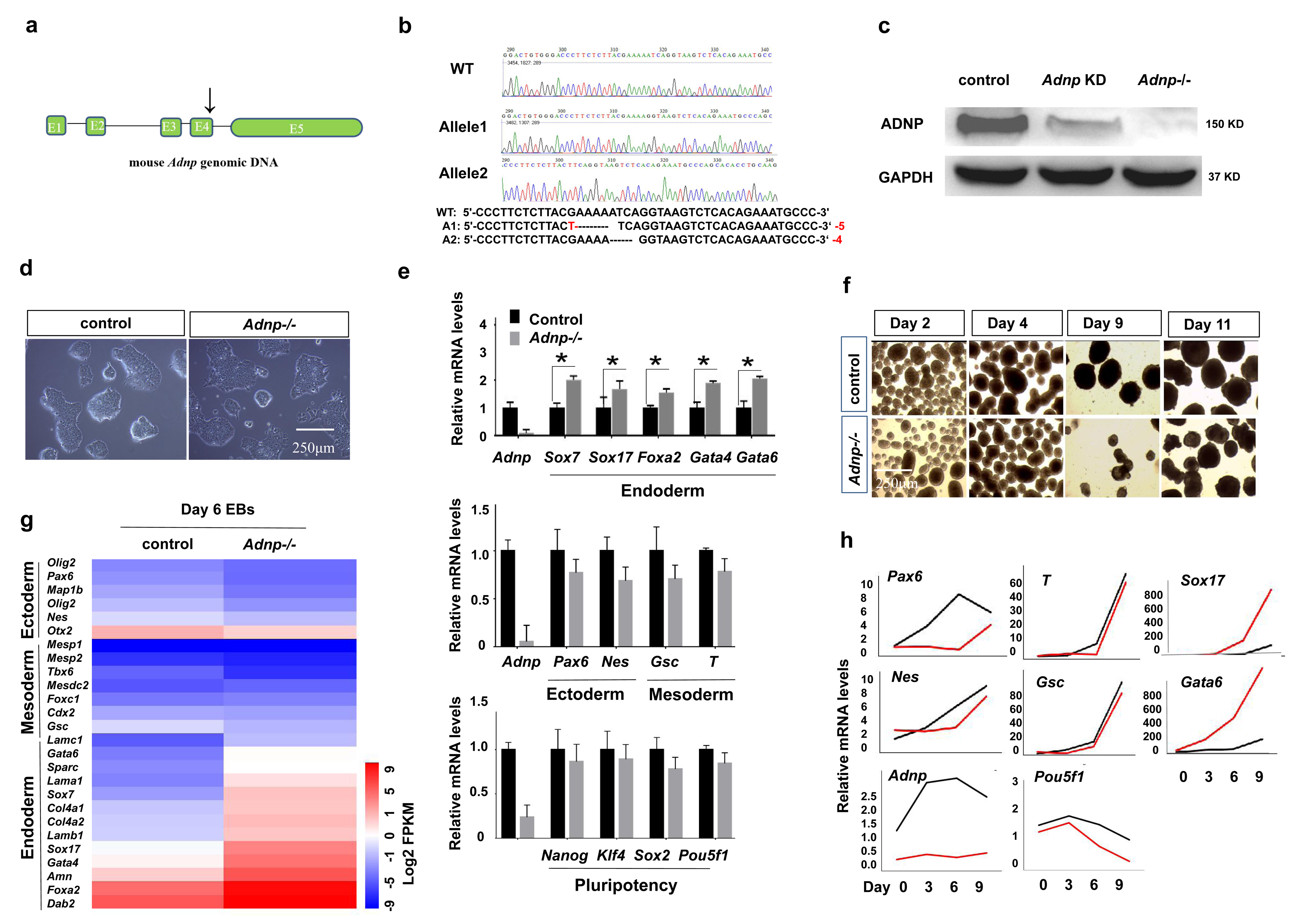
ADNP depletion leads to loss of ESC phenotype. **a** Cartoon depicting the gRNA target sites at exon 4 of mouse *Adnp* gene. **b** Genotyping showing the mutant alleles. **c** Western blot analysis of ADNP levels in control, shRNA knockdown and *Adnp-/-* ESCs. WB has been repeated for three times. **d** Representative image showing morphology of control and early passaged *Adnp-/-* ESCs. **e** The mRNA expression levels of representative pluripotency-related, mesodermal, neuroectodermal, endodermal genes in control and early passaged *Adnp-/-* ESCs. qPCR has been repeated for at least three times. **f** Representative image showing morphology of embryoid bodies (EBs) at indicated time points. **g** Heat-map of DEGs of indicated lineage-specific genes for control and *Adnp-/-* ESC-derived day 6 EBs, based on three RNA-seq replicates. **h** qPCR analysis showing the dynamic expression of lineage-specifying genes during EB formation of control and *Adnp-/-* ESCs. qPCR experiments have been done for three times.

In the traditional self-renewal medium containing LIF-KSR plus FBS, the newly established *Adnp-/-* ESC colonies overall exhibited typical ESC-like morphology (Fig. 1d). To understand how loss of *Adnp* affected the global gene expression, we performed RNA-sequencing (RNA-seq) experiments for control and early passaged *Adnp-/-* ESCs. Our RNA-seq analysis showed that the primitive endoderm genes such as *Gata6*, *Gata4*, *Sox17*, *Sox7* and *Sparc* were slightly up-regulated in the absence of ADNP. However, loss of ADNP effects on pluripotency-related, mesodermal and neuroectodermal genes were subtle. Our qRT-PCR and immunofluorescence assay confirmed the RNA-seq results (Fig. 1e; Supplementary Fig. 1a). Consistently, *Adnp-/-* ESCs can maintain self-renewal capacity for many generations before eventually adopting a flatten morphology and exhibiting reduced alkaline phosphotase activity (Supplementary Fig. 1b). Thus, our results indicated that acute ADNP depletion in ESCs does not result in sudden and complete loss of self-renewal, while prolonged ADNP depletion causes loss of ESC phenotype in the LIF/KSR medium.

To confirm that the observed phenotypes were not due to the possible off-target effect of the CRISPR/Cas9 technology, we also reduced *Adnp* transcript levels in ESCs by shRNA knockdown approach. The infection of lentivirus made from each of the two individual shRNAs led to about 70-80% reduction of *Adnp* mRNA levels (Fig. 1c; Supplementary Fig. 1c). *Adnp* knockdown ESCs in the LIF-KSR medium exhibited similar morphological characteristics and gene expression profile to *Adnp*-/- ESCs, and could be passaged for many generations without obvious morphological change (Supplementary Fig. 1d,e).

### ADNP promotes ES cell directional neural differentiation

Next, we examined the pluripotency of *Adnp-/-* ESCs by performing the classical embryoid bodies (EBs) formation assay. Control ESCs were capable of forming large smooth spheroid EBs, whereas *Adnp-/-* ESCs formed smaller rough disorganized structures (Fig. 1f). To understand how loss of *Adnp* affected the global gene expression, we performed RNA sequencing (RNA-seq) using day 6 EBs derived from control and *Adnp-/-* ESCs. A total of 2,004 differentially expressed genes (DEGs) (log2 fold change>1.5 and *p≤ 0.05*) were identified based on two RNA-seq replicates. Of which, an average of 1,088 genes were significantly up-regulated and 916 genes were down-regulated. Extra-embryonic endodermal genes such as *Gata6*, *Gata4*, *Sox17*, *Sox7* and *Foxa2* were substantially up-regulated, while pluripotency genes such as *Nanog* and *Sox2*, and neural-ectodermal genes such as *Pax6, Olig2, Sox1, Nestin* and *Wnt1* were significantly down-regulated (Fig. 1g). Our qRT-PCR assay of selected genes confirmed the RNA-seq results (Fig. 1h). Thus, based on the ESC-derived embryoid body (EB) model, ADNP is required for the neuroectodermal differentiation of mESCs, which was in line with the recent report^4^.

To better understand the role of ADNP in ESC neural differentiation, we directly differentiated control and *Adnp-/-* ESCs towards a neural cell fate. We initially used a published serum-free adherent monolayer culture protocol using N2B27 medium^9^. Control ESCs could be efficiently differentiated into neuro-ectodermal precursors and further induced into mature neurons. Unfortunately, *Adnp-/-* ESCs failed to do so as the majority of cells died in about 3-4 days (data not shown). The massive cell death in this culture system prevented us from further analyzing the role of ADNP in neural induction. To overcome this issue, we adapted a modified three-dimensional (3D) floating neurosphere protocol that allows fast yet efficient differentiation of ESCs into neural progenitors (Stage 1) and mature neurons (Stage 2) (Fig. 2a). The key of this protocol is the addition of FGF and EGF growth factors in Stage 1. It is known that FGF signaling plays a pivotal role in neural lineage specification through a “autocrine” induction mechanism^9, 10^. And the growth factor EGF and bFGF synergistically promote neural progenitor cell survival, proliferation and neuronal cell commitment^11, 12^.

**Fig. 2.**
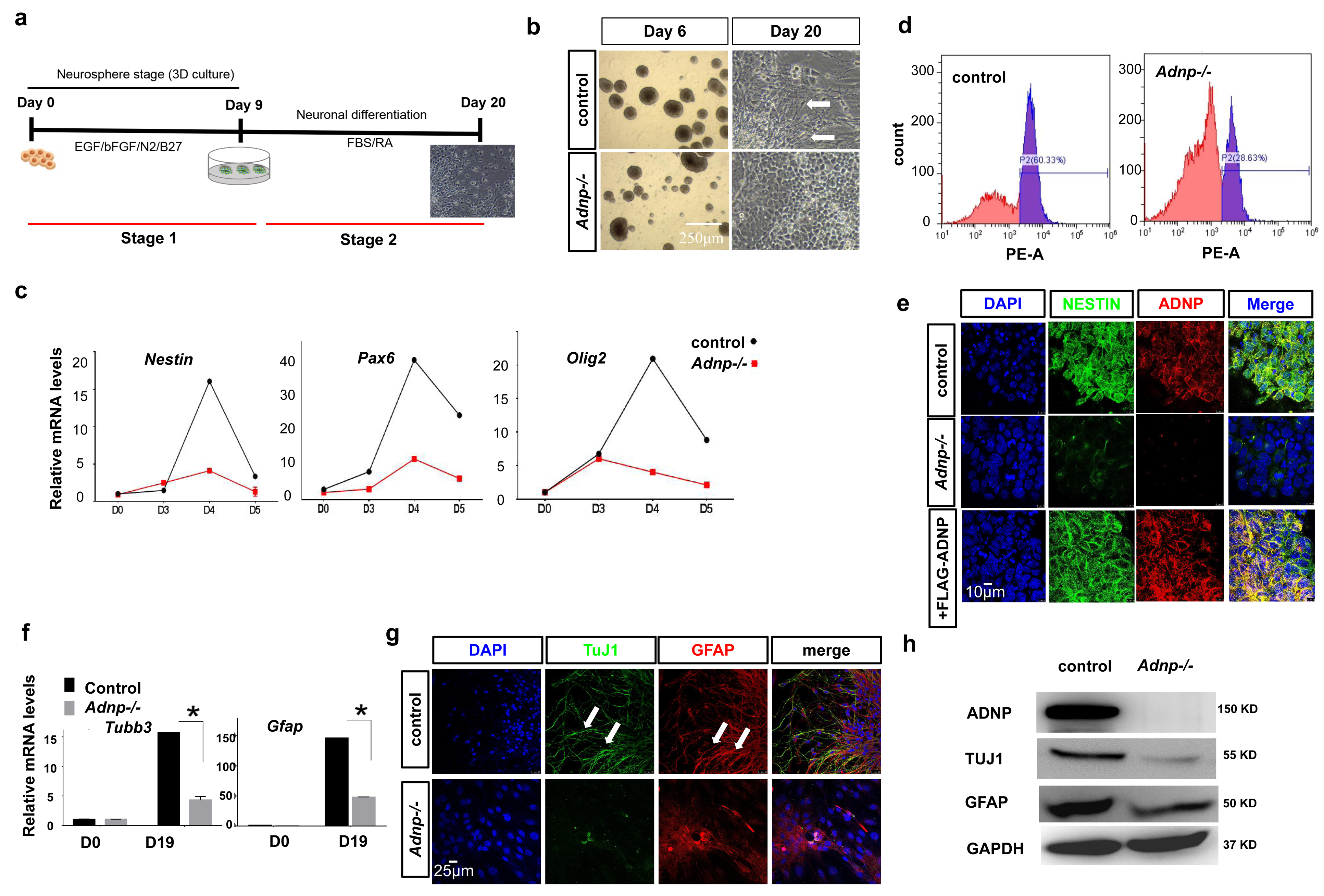
ADNP is required for proper ESC neural differentiation. **a** Cartoon showing the 2-Stage ESC neural differentiation protocol. At Stage 1, ESCs were cultured in suspension to form neurospheres with indicated growth factors; at Stage 2, the neurospheres were plated onto 0.1% gelatin-coated plates to allow further differentiation for an additional 7-14 days. **b** Representative image showing morphology of day 6 and day 20 control and *Adnp-/-* ESC-derived neurospheres neuronal cultures. The white arrow showing the fibre-like neuronal structures. **c** The dynamic expression profile of representative neuroectoderm genes during control and *Adnp-/-* ESC neural induction. Experiments were repeated for three times. **d** Flow cytometry analysis for quantification of PAX6^+^ cells. Red: isotype control; purple: experimental group using PAX6 antibody. **e** IF staining of neural progenitor markers NESTIN and ADNP for control, *Adnp-/-* ESC and FLAG-ADNP restoring *Adnp-/-* ESC-derived day 6 neurospheres. **f** qRT-PCR analysis for the expression of *Tubb3* (encoding TuJ1) and *Gfap* for day 19 control and *Adnp-/-* ESC-derived neuronal cell cultures. **g** IF staining of neuronal marker TuJ1 and glial marker GFAP for day 19 control and *Adnp-/-* ESC-derived neuronal cell cultures. The white arrows showing the neuronal fibre structures. **h** WB analysis of TuJ1 and GFAP levels in day 19 control and *Adnp-/-* ESC-derived neuronal cell types. All experiments were repeated for three times, and shown are the representative data.

At the end of Stage 1 of ESC neural differentiation, neurospheres from control ESCs were large, round and even (Fig. 2b). In contrast, neurospheres from *Adnp-/-* ESCs were smaller, rough and disorganized. The qRT-PCR assay showed that day 6 control ESC-derived neurospheres abundantly expressed *Olig2*, *Pax2*, *Pax6* and *Nestin*, indicating of efficient formation of neural progenitor cells (NPCs) (Fig. 2c; Supplementary Fig. 2a). Flow cytometric assay revealed that more than 60% of NPCs were PAX6^+^, indicating that our ESC neural induction was of high efficiency (Fig. 1d). When day 6 *Adnp-/-* ESC-derived neurospheres were analyzed, the expression of neural progenitor markers such as *Olig2*, *Pax2*, *Pax6* and *Nestin* was substantially reduced (Fig. 2c-e). Flow cytometric assay showed that only 29% of *Adnp-/-* ESC-derived NPCs were PAX6^+^. Thus, the ability of ESCs to differentiate into NPCs was severely compromised in the absence of ADNP.

When day 9 neurospheres from control ESCs were plated onto gelatin-coated plates to allow further differentiation for an additional 7-14 days (Stage 2), obvious neuronal fiber-like structures characterized by the branched and elongated morphology, were observed (Fig. 2b). qRT-PCR analysis showed that day 19 ESC-derived cells abundantly expressed neuronal and glial marker genes *Tubb3* and *Gfap*, indicating that more mature neural cell types such as neurons and astrocytic cell types were formed (Fig. 2f). IF (immunofluoresence) staining showed that TuJ1 and GFAP were abundantly expressed in these cells (Fig. 2g, h). In contrast, when neurospheres from *Adnp-/-* ESCs were plated onto gelatin-coated plates for further differentiation, few neuronal fibers were observed at day 19 (Fig. 2b). qRT-PCR, IF and WB analysis showed that the expression of GFAP and TuJ1 was substantially reduced in day 19 *Adnp-/-* ESC-derived neural cell cultures compared to control counterparts (Fig. 2f-h; Supplementary Fig. 2b, c).

Taken together, we concluded that ADNP is required to promote ESC neural differentiation, and its loss led to defective formation of neural progenitor cells and mature neuronal cell types.

### Loss of ADNP blocks neural induction by inhibiting the expression of early neuroectoderm developmental genes

During ES cell neural differentiation, ADNP levels were gradually induced, reaching its maximum level of expression at around day 4 (Supplementary Fig. 3a). The dynamic expression pattern of *Adnp* was closely correlated to that of the key neuroectoderm developmental genes such as *Olig2*, *Pax6* and *Nestin* (Supplementary Fig. 3b). Importantly, loss of ADNP caused a significant down-regulation of these genes (Fig. 2c). These observations implied that ADNP may function at early stage of neural differentiation program by controlling the expression of early neuroectoderm developmental genes.

To gain insights into the ADNP-dependent transcriptional program during ES cell differentiation toward a neural cell fate, we monitored the dynamic changes of gene expression at different time points of neural induction, by performing RNA-seq experiments at day 0, day 3, and day 6 of neural differentiation from control and *Adnp-/-* ESCs (Fig. 3a). In day 3 ESC-derived neurospheres, ∼1,200 DEGs (log2 fold change>1.5 and *p≤ 0.05*) were identified, of which 627 genes were significantly up-regulated, and 578 genes were significantly down-regulated in the absence of ADNP. Among the down-regulated genes were enriched for neuroectodermal markers such as *Sox3*, *Fgf5*, *Foxd3*, *Nestin*, *Sall2* and *Nptx2,* while the up-regulated genes were enriched for pluripotency-related markers such as *Dppa4*, *Esrrb*, *Klf4*, *Sox2* and *Zfp57*, as well as primitive endodermal markers such as *Sox7*, *Gata6, Gata4, ApoE, Fabp3*, *Cubn* and *Cited*. In day 6 ESC-derived neural progenitors, ∼2,430 DEGs (log2 fold change>1.5 and *p≤ 0.05*) were identified, of which 1,344 genes were significantly up-regulated, and 1,086 genes were significantly down-regulated (Fig. 3b). Of note, genes implicated in neuroectoderm development such as *Fgf5*, *Sox1*, *Sox3*, *Pax2/6*, *Pax3/7*, *Foxd3*, *Nestin*, *Irx2*, *Cdx2*, *Sall2* and *Nptx2* were significantly down-regulated. Heat map analysis of day 3 and day 6 DEGs showed that there was a significantly reduced expression of neuroectoderm-related genes, and a significantly increased expression of pluripotency-related genes in the absence of ADNP (Fig. 3c; Supplementary Fig. 3c). The qRT-PCR assay confirmed the RNA-seq results (Fig. 2c). These data strongly suggested that ADNP performs an important role at the early stage of neural differentiation by promoting the expression of neuroectoderm developmental genes and inhibiting pluripotency-related genes.

**Fig. 3.**
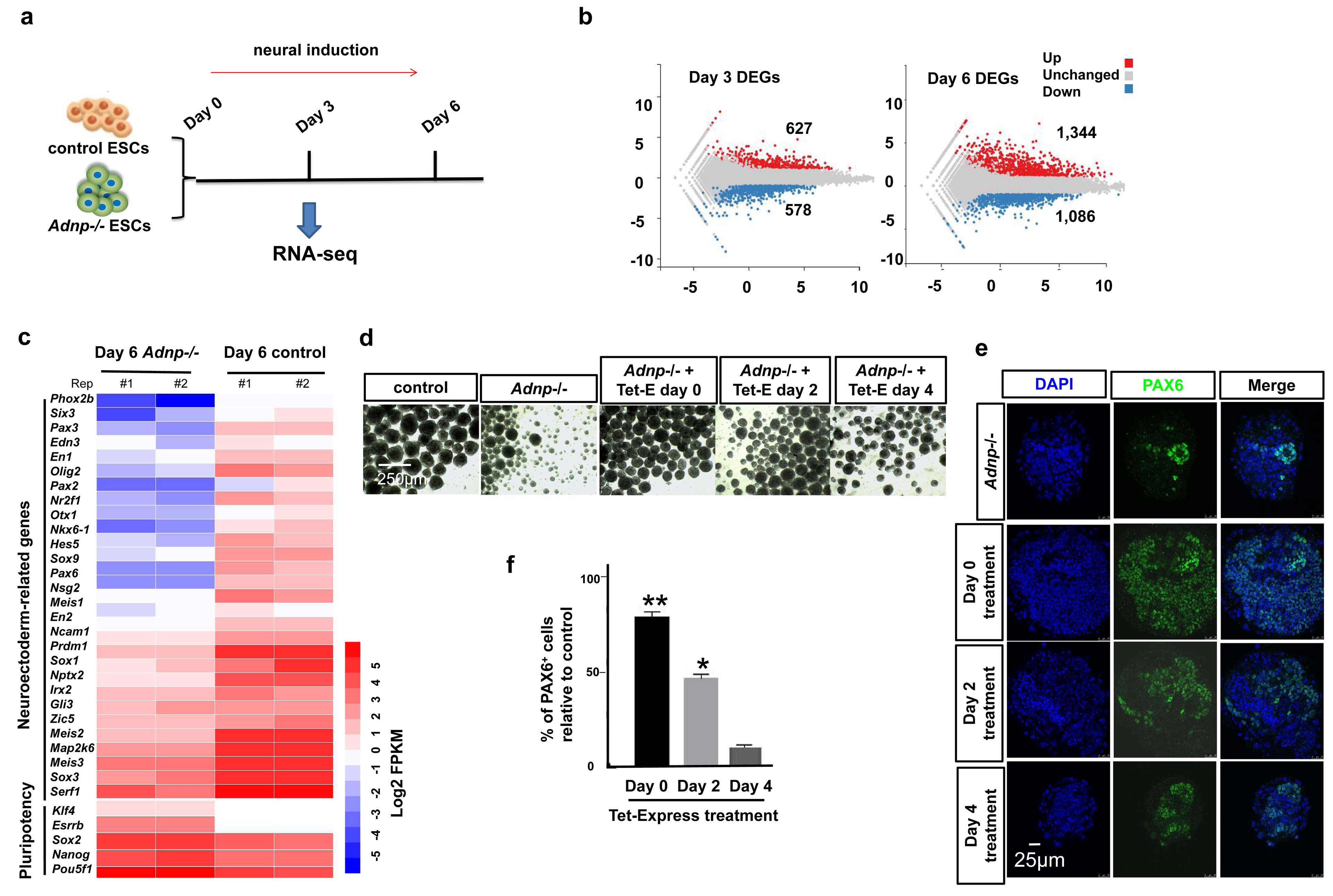
ADNP promotes the expression of neuroectoderm developmental genes. **a** Schematic representation of design of RNA-seq experiments; **b** Plot showing DEGs from day 3 and day 6 control and *Adnp-/-* ESC-derived neurospheres; RNA-seq for each time points were repeated for at least two times. The numbers of up- and down-regulated genes were shown as average. **c** Heatmap illustrating the expression of selected neuroectoderm and pluripotency-related genes that were shown as log2 FPKM in day 6 control and *Adnp-/-* ESC-derived neurospheres. Each lane corresponds to an independent biological RNA-seq sample. **d** Representative morphology of day 6 *Adnp-/-* ESC-derived neurospheres after addition of Tet-Express proteins from the indicated time points. Tet stands for Tet-Express. Experiments were repeated for two times. **e** IF staining of PAX6 for day 6 *Adnp-/-* ESC-derived neurospheres after addition of Tet-Express proteins at the indicated time points. **f** Quantitative analysis of PAX6^+^ cells based on panel **(e)**. All IF staining experiments were repeated for at least two times, and shown are the representative data.

If ADNP functions upstream of the key neuroectoderm developmental genes to control neural induction, restoring ADNP should rescue the neural differentiation defects from *Adnp-/-* ESCs. To this end, we took use of a Tet-Express inducible transgenic *Adnp-/-* ES cell line in which 3×FLAG-ADNP can be induced by the addition of Tet-Express protein (Supplementary Fig. 3d). We added Tet-Express at different time points during mutant ES cell neural induction: day 0, day 2, and day 4, and collected day 6-7 ESC-derived neurospheres for further analysis. We found that the addition of Tet-Express at early time points (day 0 or 2) could partially rescue the morphological and gene expression defects, while addition of Tet-Express after day 4 had minimal effects (Fig. 3d-f). When ESC-derived neurospheres were allowed for further differentiation, day 19 cell cultures treated with Tet-Express at early time points showed no significant differences from control cultures. By contrast, cultures treated with Tet-Express at or after day 4 exhibited few neuronal fiber-like structures (Supplementary Fig. 3e).

Based on the above data, we concluded that ADNP plays important roles at the early stages of ESC neural differentiation by promoting the expression of key developmental genes of neuroectodermal lineage.

### Wnt/β-catenin signaling is reduced in the absence of ADNP

Next, we sought to explore the underlying mechanisms by which ADNP controls ES cell neural induction and neuronal differentiation. To this end, we performed KEGG pathway enrichment analysis of differentially expressed genes (DEGs) identified in our RNA-sequencing results. This analysis revealed a strong enrichment among DEGs for annotations associated with signal transduction as well as signaling molecules and interaction (Supplementary Fig. 4a). Detailed analysis showed that ADNP-dependent down-regulated genes were highly enriched for gene networks regulating Wnt signaling (Fig. 4a). Consistently, heat map analysis of DEGs revealed that there was a significant reduction in transcript levels of Wnt pathway-related genes (Fig. 4b; Supplementary Fig. 4b). This observation strongly suggested that loss of ADNP blocks Wnt signaling pathway. To confirm the RNA-seq results, qRT-PCR for day 3 and day 9 ESC-derived neurospheres was performed to analyze the expression of Wnt target genes such as *Ccnd1, Axin2, Lef1* and *Tcf712.* The expression of these genes at the indicated time points was greatly reduced in the absence of ADNP (Fig. 4c). Note that the expression of *Ctnnb1* gene, which encodes for β-Catenin, was barely changed in the absence of ADNP. To further confirm that Wnt signaling was compromised in the absence of ADNP, we performed TopFlash luciferase reporter assay for day 3 control and mutant ESC-derived neural progenitor cells. As shown in Fig. 4d, loss of ADNP substantially decreased the TopFlash reporter activities, in the absence and presence of Wnt3a.

**Fig. 4.**
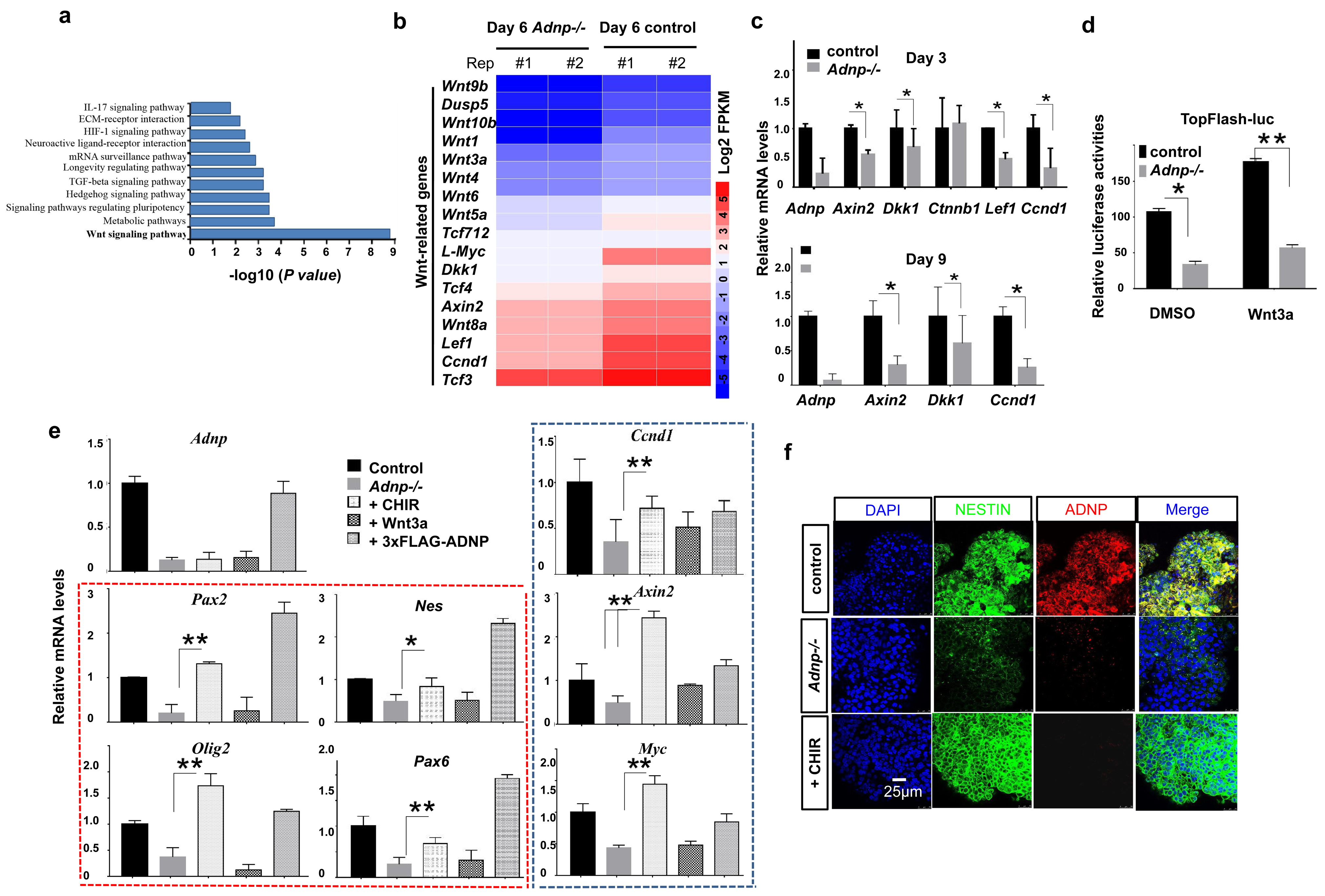
Wnt signaling is impaired in the absence of ADNP. **a** Dissection of KEGG data of DEGs from day 3 and day 6 control and *Adnp-/-* ESC-derived neurospheres, showing enrichment of the Wnt signaling pathway. **b** Heatmap illustrating the expression of selected Wnt-related genes that were shown as log2 FPKM in day 6 control and *Adnp-/-* ESC-derived neurospheres. Each lane corresponds to an independent biological RNA-seq sample. **c** qRT-PCR analysis for the indicated Wnt target genes for day3 and day9 control and *Adnp-/-* ESC-derived neurospheres. The experiments were repeated for three times. **d** Top-Flash luciferase activity assay for lysates from day 3 control and *Adnp-/-* ESC-derived neurospheres, in the absence or presence of Wnt3a. Experiments were repeated for two times. **e** Rescue of the expression of the indicated neural developmental genes and putative Wnt target genes by addition of CHIR and Wnt3a, or by restoring 3×FLAG-ADNP. Genes in red box are representative neural developmental genes, and genes in blue box are representative Wnt target genes. The experiments were repeated for three times. **f** Rescue of NESTIN expression by addition of CHIR. Representative IF staining of NESTIN for day 6 control and *Adnp-/-* ESC-derived neurospheres.

On the other hand, we investigated whether overexpressing ADNP could facilitate Wnt signaling. To this end, we performed the TopFlash reporter assay in HEK293T cells transfected with an increasing amount of plasmids encoding ADNP. As shown in Supplementary Fig. 4c, the TopFlash reporter activities were gradually enhanced in a ADNP dose-dependent manner, in the presence and absence of Wnt3a. Based on these data, we concluded that ADNP positively regulates Wnt/β-catenin signaling.

Wnt signaling is well-known for its critical role in neural cell induction, proliferation and differentiation, and dysfunction of this signaling pathway is frequently associated with neural developmental disorders^13^. The above data suggested that loss of ADNP function may affect neural induction by inhibiting Wnt signaling. If this was the case, enhancing Wnt signaling should rescue the neural induction defects from *Adnp-/-* ESCs. We therefore performed rescue experiments by the addition of CHIR99021 or WNT3a from early time points (from day 0 or 1) of mutant ESC neural differentiation. CHIR99021 (CHIR), a potent and highly selective inhibitor of glycogen synthase kinase 3β (GSK3β), is a well-characterized canonical Wnt signaling pathway activator. We have tested different concentration of CHIR99021, and found that the addition of 3 μM CHIR99021 partially rescued the gene and protein expression as well as morphological defects of day 6 *Adnp-/-* ESC-derived neurospheres (Fig. 4e, f; Supplementary Fig. 4d, e). However, addition of different doses of WNT3a proteins had little effect on the gene expression profiles. This data suggested that ADNP may regulate Wnt signaling downstream of the activity of the Wnt ligands but upstream or parallel to GSK3β which is known to phosphorylate and induce proteasomal degradation of β-Catenin^14^.

Taken together, we concluded that ADNP is critical for ES cell neural differentiation by promoting the Wnt/β-catenin signaling pathway. In the absence of ADNP, Wnt signaling was impaired and the ability of ESC differentiation towards neuroectodermal lineage cells was compromised.

### Identifying β-Catenin as an ADNP interacting protein

In our immunoprecipitation combined with mass spectrometry assay, β-Catenin was detected as one of the top hits of ADNP immunoprecipitates (Fig. 5a). As β-Catenin is the key player in the canonical Wnt signaling pathway, we hypothesized that ADNP may regulate Wnt signaling by targeting β-Catenin during ESC differentiation towards a neural fate. We first investigated whether ADNP interacts with β-Catenin by performing co-immunoprecipitation (Co-IP) experiments. The Co-IP results showed that endogenous ADNP can interact with β-Catenin in both ESCs and day 3 ESC-derived neurospheres, exhibiting much stronger interaction in the neurospheres (Fig. 5b, c). We also performed the Co-IP experiments using the 3×FLAG-*Adnp* overexpressing *Adnp-/-* ESC line. As shown from Figure 5d, FLAG-ADNP readily pulled down endogenous β-Catenin in day 3 ESC-derived neural progenitors (Fig. 5d).

**Fig. 5.**
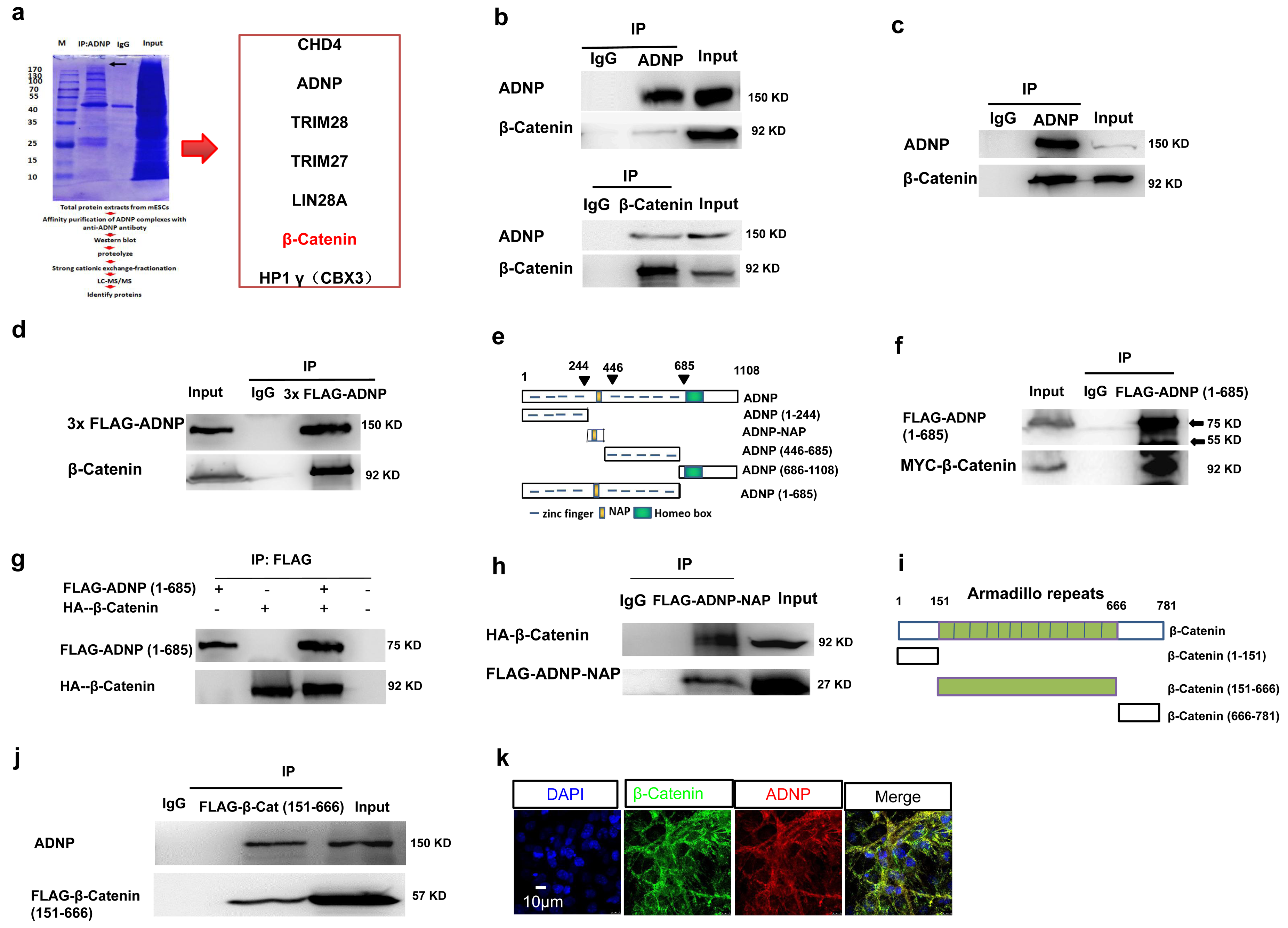
Identifying β-Catenin as a ADNP interacting protein. **a** Schematic representation showing the experimental design of IP in combination mass spectrometry assay (left) and a list of representative ADNP interacting proteins (right). **b** Co-IP data for endogenous ADNP and β-Catenin in ESCs. Up: IP ADNP followed by WB β-Catenin; bottom: IP β-Catenin followed by WB ADNP. **c** Co-IP data for endogenous ADNP and β-Catenin of day 3 ESC-derived neurospheres. **d** Co-IP data for FLAG-ADNP and β-Catenin in 3×FLAG-ADNP overexpressing *Adnp-/-* ESCs. **e** Schematic representation of the full-length and the truncated ADNP mutants. **f** IP of FLAG-ADNP-Nter (1-685) and MYC-β-Catenin in 293T cells. **g** IP of in vitro synthesized HA-β-Catenin and FLAG-ADNP. **h** Co-IP of FLAG-ADNP-NAP and HA-β-Catenin in 293T cells. **i** Schematic representation of the full-length and the truncated β-Catenin mutants. **j** Co-IP of FLAG-β-Catenin (151-666) and ADNP in 293T cells. **k** Colocalization of ADNP and β-Catenin in day 19 ESC-derived neuronal cell cultures. All WB experiments have been repeated for three times, and shown are the representative images.

Next, we mapped the interacting domains for ADNP and β-Catenin by co-transfecting plasmids encoding the truncated form of ADNPs and β-Catenin into 293T cells (Fig. 5e). We found that it was the N-terminal fragment (1-685) but not the C-terminal fragment of ADNP (686-1108) that is responsible to interact with β-Catenin (Fig. 5f; Supplementary Fig. 5a). To investigate whether they interact directly, we used a reticulate system to synthesize FLAG-ADNP (1-685) and HA-β-Catenin. When they were mixed together, FLAG-ADNP (1-685) could readily pull down HA-β-Catenin (Fig. 5g). Besides the Zinc fingers, ADNP-Nter (1-685) contains NAP (which is also called davunetide or CP201), an 8-amino acid neuroprotective peptide (NAPVSIPQ) (354-361 within the ADNP-Nter) derived from ADNP protein^15^. It was shown that NAP could enhance microtubule assembly and neurite outgrowth by interacting with tubulin, and protect neurons and glia exposed to toxins in mouse and rat models^15, 16^. Recently, it was shown that the major symptoms of the Helsmoortel-Van der Aa syndrome, such as developmental delays, vocalization impairments, motor and pace dysfunctions, and social and object memory impediments, could be partially ameliorated by daily NAP administration^17, 18^. We wondered whether NAP alone could be sufficient to interact with β-Catenin. When we transfected plasmids encoding FLAG-ADNP-NAP and HA-β-Catenin into 293T cells, FLAG antibodies could pull down HA-β-Catenin (Fig. 5h). Of note, ADNP-Nter without NAP could still interact with β-Catenin but this interaction was much weaker than that of ADNP-Nter and β-Catenin (Supplementary Fig. 5b).

Next, we determined the fragments of β-Catenin that are responsible for interaction with ADNP. β-Catenin has an N-terminal domain (residues 1-150) that harbors the binding site for GSK3β and CK1 phosphorylation, a central armadillo domain (residues 151-666) composed of 12 armadillo repeats, and a C-terminal domain (residues 667-781) (Fig. 5i). Our mapping experiments showed that the central armadillo domain of β-Catenin strongly interacts with ADNP, while the N-terminal (1-150) and the C-terminal fragment (667-781) barely interact with ADNP (Fig. 5j; Supplementary Fig. 5c). These data indicated that the armadillo domain of β-Catenin primarily mediates the interaction with ADNP.

Finally, we performed IF experiments to investigate the cellular co-localization of ADNP and β-Catenin. During ES cell neuronal differentiation, there was a marked change in the intra-cellular localization of ADNP proteins: from predominantly nuclear localization in undifferentiated ESCs to predominantly cytoplasmic localization in ESC-derived NPCs and mature neurons (Supplementary Fig. 5d). In ESC-derived NPCs and mature neurons, ADNP was extensively co-localized with β-Catenin in the cytosol (Fig. 5k).

Taken together, we identified β-Catenin as an ADNP interacting protein, which suggested that ADNP had important roles in the regulation of Wnt/β-catenin signaling by physical association with cytosolic β-Catenin.

### ADNP stabilizes β-Catenin during ESC neural differentiation

Given the physical association of ADNP and β-Catenin, we asked whether ADNP is involved in the turnover of β-Catenin protein. First, we examined the expression levels of β-Catenin in day 3 control and *Adnp-/-* ESC-derived neurospheres. Loss of ADNP led to a significant reduction of total β-Catenin levels (Fig. 6a). This reduction was primarily contributed by the alteration of the cytoplasmic fraction of β-Catenin. In fact, the levels of the nuclear fraction of β-Catenin were barely changed in the absence of ADNP (Fig. 6b). Throughout ESC neural differentiation, total β-Catenin levels remained lower in the absence of ADNP than its presence (Fig. 6c, d; Supplementary Fig. 6a, b). These observations strongly suggested that ADNP regulates the turnover of β-Catenin which primarily occurred in the cytosol. When ESC-derived neural progenitors were treated with the proteasome inhibitor MG132, loss of ADNP failed to induce β-Catenin degradation, reflecting that ADNP-regulated β-Catenin degradation was dependent on the ubiquitin-proteasome pathway (Fig. 6e). Consistently, loss of ADNP strongly enhanced the ubiquitylation levels of β-Catenin, which could be reversed by restoring 3×FLAG-ADNP in *Adnp-/-* ESC-derived neurospheres (Fig. 6f; Supplementary Fig. 6c).

**Fig. 6.**
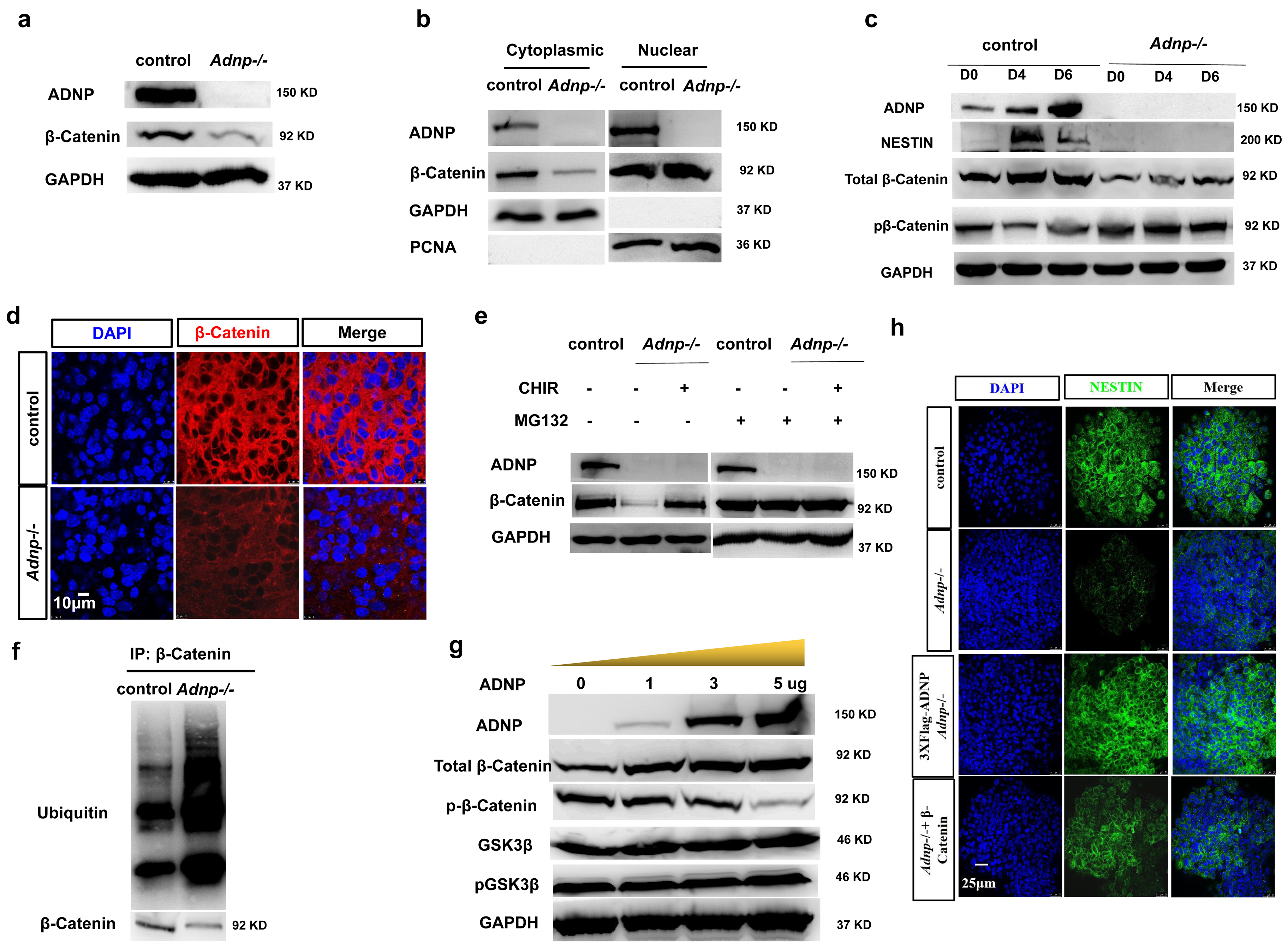
ADNP stabilizes β-Catenin during ESC neural differentiation. **a** WB showing total β-Catenin levels in day 3 control and *Adnp-/-* ESC-derived neurospheres. **b** WB showing β-Catenin levels in the cytoplasmic and the nuclear fraction of lysates from day 3 control and *Adnp-/-* ESC-derived neurospheres. Note that β-Catenin levels were reduced in the cytoplasmic but not in the nuclear fraction of lysates. **c** Time course WB assay showing total and phosphorylated β-Catenin levels during control and mutant ESC differentiation towards a neural fate. **d** Representative IF staining of β-Catenin in day 19 ESC-derived neuronal cultures; **e** Representative WB showing β-Catenin levels in the combination treatment of CHIR and MG132. **f** Representative WB showing the ubiquitylation levels of β-Catenin in control and *Adnp-/-* ESCs. **g** Representative WB showing the expression of the indicated proteins in 293T cells that transfected with an increasing dose of ADNP. **h** Representative IF staining of NESTIN showing the rescue of NESTIN in day 6 Tet-Express inducible HA-β-catenin *Adnp-/-* ESC derived neurospheres by the addition of Tet-Express every day at the early stage of neural induction.

On the other hand, we investigated the effect of overexpressing ADNP on total β-Catenin levels in HEK293T cells. Overexpressing ADNP led to higher expression of β-Catenin (Fig. 6g). In addition, overexpressing ADNP increased β-Catenin levels in a dose dependent fashion. Next, we asked whether overexpression of ADNP proteins affects the stability of coexpressed β-Catenin by co-transfecting plasmids encoding FLAG-ADNP and HA-β-Catenin into 293T cells. We found that FLAG-ADNP could stabilize HA-β-Catenin in a dose-dependent fashion (Supplementary Fig. 6d). Combining both loss of function and gain of function data, we concluded that ADNP is required for the stability of β-Catenin in ESCs and during ESC differentiation towards a neural cell fate.

It is well-known that β-Catenin is subjected to phosphorylation by GSK3β who destabilizes β-Catenin by phosphorylating it at Ser33, Ser37 and Thr41. A previous study reported that there might be an increase in the ratio of active/inactive GSK3β in *Adnp+/-* mice, suggesting that ADNP may negatively regulate the phosphorylation activity of GSK3β^8^. To investigate whether ADNP regulates the stability of β-Catenin through modulating GSK-3β activities, WB analysis for phospho-GSK3β (Ser9) and total GSK3β were performed for day 3 control and *Adnp-/-* ESC-derived neurospheres. We found that both the phospho-GSK3β and total GSK3β levels were barely altered in the absence of ADNP (Supplementary Fig. 6e). In addition, overexpression of ADNP had little effect on phospho-GSK3β and total GSK3β levels (Fig. 6g). These results suggested that ADNP does not modulate the levels or activities of GSK3β.

Next, we asked whether ADNP is involved in the regulation of β-Catenin phosphorylation by GSK3β. We found that this was the case. As shown in Fig. 6a-g, phospho-β-Catenin levels were elevated in the absence of ADNP and were significantly reduced in ADNP overexpressing cells. To understand the molecular mechanism by which ADNP may regulate β-Catenin phosphorylation levels, we further mapped the armadillo repeat domain that is responsible for interacting with ADNP. The armadillo repeat domain of β-Catenin has 12 armadillo repeats, containing overlapping binding sites for many binding partners, including E-cadherin, TCF, Axin and APC^14^. It is well-known that Axin, APC and β-Catenin form the degradation complex that promotes the GSK3β-mediated phosphorylation and subsequent degradation of β-Catenin. We divided the armadillo repeat domain into two parts: one contains repeats 1-4 (Arm1) which are known to mediate β-Catenin for the interaction with the scaffolding protein Axin, and another one composed of repeats 5-12 (Arm2) which are known to mediate β-Catenin for the interaction with APC (Fig. 7a)^19^. By performing IP experiments, we found that both the Arm1 and Arm2 were able to interact with ADNP although Arm2 displayed a stronger capacity to associate with ADNP (Fig. 7b, c). Based on the the result, we hypothesized that ADNP may protect β-Catenin from hyperphosphorylation by GSK3β through competing with both components of the degradation complex: Axin and APC. To directly investigate whether ADNP competes with Axin or APC for β-Catenin, we examined the relationship between ADNP – β-Catenin and β-Catenin – Axin interactions by performing competitive protein-binding assay. The amount of Axin co-immunoprecipitated with β-Catenin became reduced by the increased addition of ADNP (Fig. 7d). Similar result was obtained for the relationship between ADNP– β-Catenin and β-Catenin – APC interactions. The amount of APC co-immunoprecipitated with β-Catenin became reduced by the increased addition of ADNP (Fig. 7e).

**Fig. 7.**
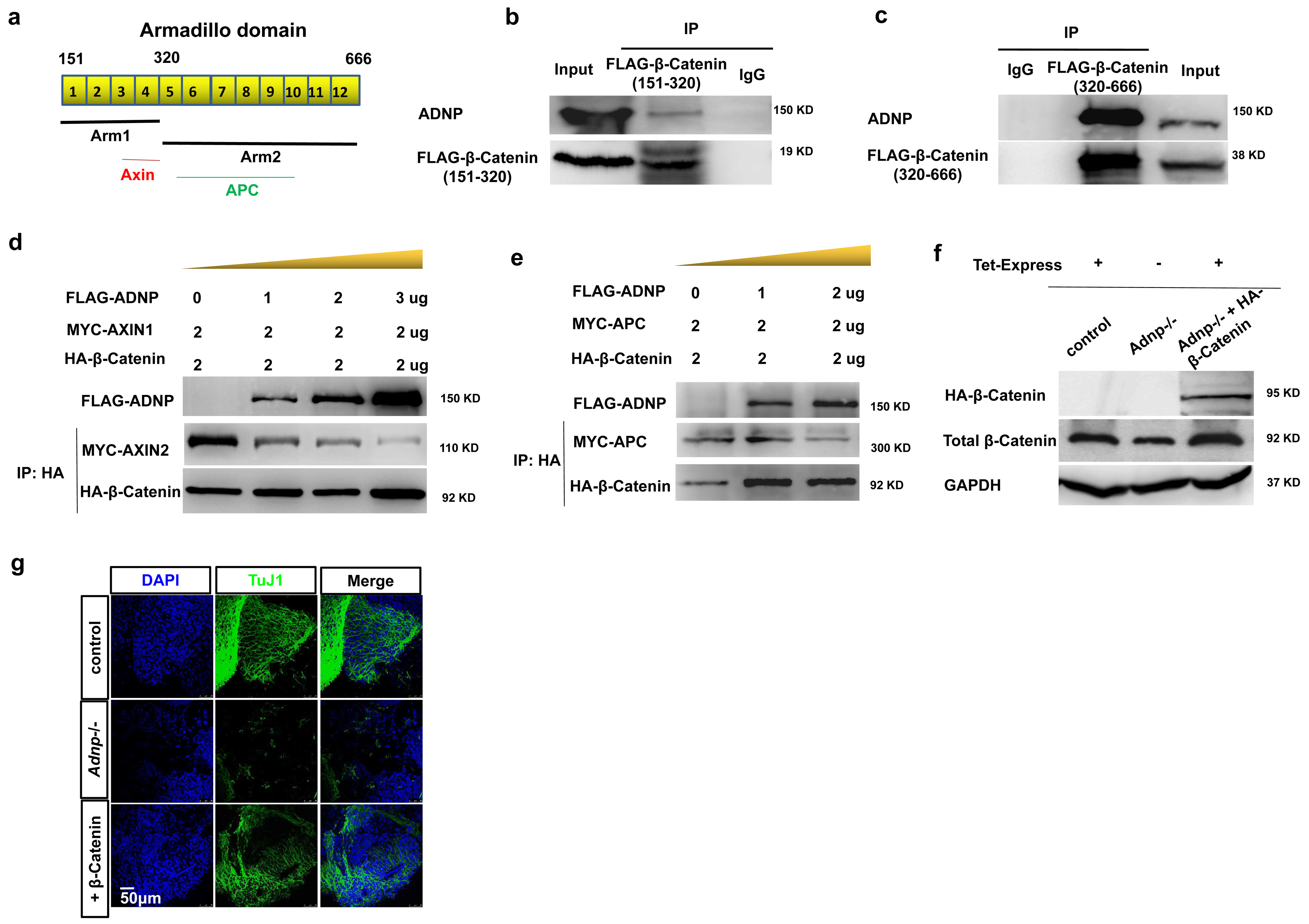
ADNP stabilizes β-Catenin by preventing the formation of degradation complex. **a** Cartoon showing the armadillo domain of β-Catenin. Arm1 contains armadillo repeat 1-4 and Arm2 contains repeat 5-9. Known core binding armadillo repeats of β-Catenin for Axin and APC were indicated with red and green bars, respectively. **b** IP experiment showing that the Arm1 fragment can interact with ADNP in 293T cells. **c** IP experiment showing that the Arm2 fragment can interact with ADNP in 293T cells. **d** The relationship between ADNP – β-Catenin and β-Catenin – Axin interactions were examined by performing competitive protein-binding assay. An increasing dose of plasmids encoding FLAG-ADNP and plasmids encoding MYC-AXIN1 and HA-β-Catenin were co-transfected into 293T cells. Lysates were extracted from the transfected cells and pull-downed by HA antibody followed by WB using MYC and HA antibodies. WB showing that the amount of Axin co-immunoprecipitated with β-Catenin became reduced by the increased addition of ADNP. **e** The competitive protein-binding assay showing that the amount of APC co-immunoprecipitated with β-Catenin became reduced by the increased addition of ADNP. **f** WB showing the expression of HA-β-Catenin by the addition of the Tet-Express in the inducible HA-β-Catenin *Adnp-/-* ESCs. **g** IF staining showing the rescue of TuJ1 signal in day 19 *Adnp-/-* ESC-derived neuronal cell cultures by adding Tet-Express transactivitor every day at early stage of neural induction. All WB experiments were repeated for at least two times.

To further confirm that ADNP can protect β-Catenin from hyperphosphorylation by GSK3β, we transfected an increasing dose of plasmids encoding wild type ADNP or ADNP-Cter (686-1108) mutants into 293T cells, and examined the effects on total β-Catenin and p-β-Catenin levels. The ADNP-Cter here served as a negative control as it was unable to interact with β-Catenin. Overexpressing wild type ADNP decreased the p-β-Catenin levels and increased the total β-Catenin levels, in a dose dependent fashion (Fig. 6g). However, ADNP-Cter overexpression had no effects on either β-Catenin or p-β-Catenin levels (data not shown). This data suggested that the interaction between ADNP and β-Catenin was required for the ADNP mediated stabilization of β-Catenin through modulating its phosphorylation levels.

Finally, we asked whether increasing β-Catenin protein levels could rescue the neural induction defects caused by loss of ADNP. To this end, we made a Tet-Express inducible HA-β-catenin transgenic *Adnp-/-* ESC line and used it for neural induction. Before performing rescue experiments, we confirmed that addition of Tet-Express proteins could induce the expression of total β-Catenin at similar levels to that of control ESCs (Supplementary Fig. 7f). When the Tet-Express transactivitor was added every day at the early stage of neural induction, the transgenic *Adnp-/-* ESCs could form neural progenitors and neuronal cells with gene expression profiles similar to that of wild type ESCs (Fig. 6h, 7g). Thus, restoring β-Catenin levels had similar rescue effects as CHIR99021 treatment or restoring ADNP on *Adnp-/-* ESC neural induction and differentiation. Taken together, we concluded that ADNP controls ESC neural induction primarily by stabilizing β-Catenin.

### Loss of ADNP leads to defective cerebellar neuron formation

By biochemical analysis, it has been shown that ADNP is abundantly expressed in the hippocampus, cerebral cortex and cerebellum of human and mouse brain^20, 21^. However, the spatiotemporal expression patterns of ANDP in mammalian embryonic brain have not been reported in detail. To explore this, vector slices from ten-week-old mouse embryonic brains were subjected to immunohistochemistry assay using ADNP antibodies. ADNP was expressed broadly in the embryonic brain, including olfactory bulb, cerebellum, hippocampus and cortex, with cerebellar region displaying very strong signal (Supplementary Fig. 7a). This observation suggested that ADNP is cell-autonomously required for mouse embryonic brain development and maturation, including the cerebellum. This was in line with the observation that the cerebellum and areas related to it are one of the most consistently abnormal brain regions in various neural developmental disorders^22, 23^. In fact, children with ASD or the Helsmoortel-Van der Aa syndrome often show different extent of intellectual disability and cognitive deficits, reflecting the dysfunction of the cerebellum.

We hypothesized that loss of *Adnp* may cause developmental and functional defects of embryonic cerebellum. To test this hypothesis, we took use of ESC-based in vitro cerebellar neuron differentiation as a model system, and adapted the published SFEB protocols that have been shown to efficiently induce ESCs into GABA receptor alpha 6^+^ granule cells and L7^+^ Purkinje neurons via *Math1* and *Ncam* expressing precursors^24–27^(Fig. 8a). And control and mutant ESCs were differentiated into cerebellar neurons and comparison assay was performed to gain insights into the role of ADNP in this process.

**Fig. 8.**
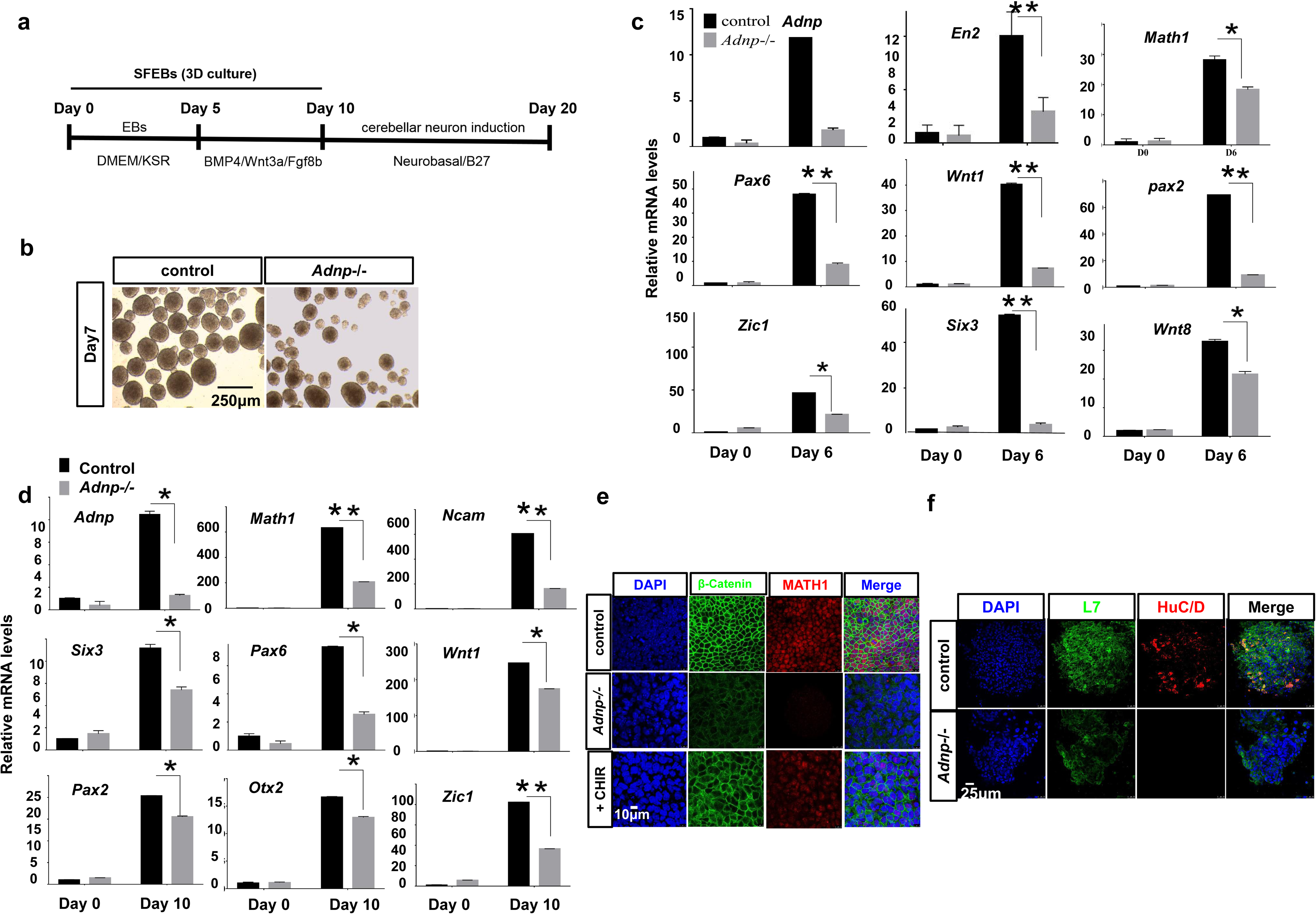
ADNP plays key role in cerebellar neuron development. **a** Schematic method for ESC differentiation into cerebellar neurons. **b** Representative morphology of day 7 control and *Adnp-/-* ESC-derived neurospheres. **c** qPCR showing the expression of neural developmental genes of day 6 control and *Adnp-/-* ESC-derived neurospheres. **d** qPCR showing the expression of EGL precursor markers of day 10 control and *Adnp-/-* ESC-derived EGL precursors. **e** IF double-staining of MATH1 and β-Catenin for day 10 control and *Adnp-/-* ESC-derived EGL precursors. **f** IF double-staining of L7 and HuC/D for day 20 control and *Adnp-/-* ESC-derived cerebellar neurons. All experiments were repeated for 3 times.

Round and large floating aggregates were formed after 5-10 days of neural differentiation from control ESCs. In contrast, the aggregates derived from *Adnp-/-* ESCs were much smaller and uneven, suggesting of defective formation of neural precursors (Fig. 8b). It has been shown that a significant proportion of SFEB/BMP4/Wnt3a/FGF8-treated ES cells at day 6 will become the external granule layer (EGL) precursors expressing *En2*, *Pax6*, *Six3* and *Zic1*^27^. We therefore examined the expression of these markers for day 6 control and *Adnp-/-* ESC-derived SFEB cultures. In control ESC-derived aggregates, *En2*, *Six3*, *Pax2*, *Zic1*, *Wnt1* and *Pax6* were abundantly expressed, suggesting that EGL precursors were successfully induced. In *Adnp-/-* ESC-derived aggregates, however, the expression of these genes was greatly reduced (Fig. 8c). After 10 days of ESC neural differentiation, aggregates derived from control ESCs highly expressed *Math1*, *Zic1* and *Ncam*, while the counterparts from mutant ESCs exhibited much lower levels of these markers (Fig. 8d, e; Supplementary Fig. 7b-d). Importantly, the reduced expression of the EGL markers in the absence of ADNP was accompanied by the reduction of β-Catenin levels (Fig. 8e). This suggested that Wnt/β-catenin signaling under the control of ADNP is required for proper generation of cerebellar neuron precursors.

The *Math1* or *Zic1* positive precursors could be further induced into GABA receptor alpha 6 positive granule cells and L7 positive Purkinje neurons on day 20 of ESC in vitro differentiation^24, 27^. The expression of Purkinje cell marker L7 was detected by immunofluorescence staining of day 20 neuronal cell cultures. As shown in Fig. 8f, day 20 aggregates from control ESCs highly expressed Purkinje cell marker L7, while the counterparts from *Adnp-/-* ESCs expressed this marker at much lower levels.

Taken together, we concluded that ADNP is required for proper cerebellar neuron formation likely by modulating Wnt/β-catenin signaling. Based on this result, we tentatively propose that cerebellar developmental defects may be important pathology of the Helsmoortel-Van der Aa syndrome.

### ADNP regulates ESC-derived neural progenitor proliferation by maintaining Wnt signaling

*Adnp-/-* embryos die in uteri displaying a striking disintegration phenotype, likely due to the marked growth inhibition commencing at about E8.0^5^. In addition, *Adnp+/−* mice show a significant neurodegeneration phenotype with reduced neuronal survival ^8^. These observations suggested that ADNP may regulate neural progenitor cell proliferation or apoptosis in a mouse model. We hypothesize that defective Wnt signaling caused by loss of function of ADNP contributes to this phenotype.

During ESC differentiation into neurospheres, a marked reduction of total neural cells was observed in the absence of ADNP (Fig. 1b, 3d). This could be confirmed by comparing the cell pellet from day 6 control and *Adnp-/-* ESC-derived aggregates (Fig. 9a). We immunostained day 6 neurospheres with a phospho-histone H3 (PH3) antibody, which is an M-phase marker, and quantitatively scored the number of proliferating cells. We found that PH3 signal in *Adnp-/-* ESC-derived aggregates was dramatically decreased compared to control ESC-derived neurospheres, suggesting that loss of ADNP led to cell proliferation defects (Fig. 9b, c). To confirm this, WB analysis was performed during neural induction of control and *Adnp-/-* ESCs using PH3 antibodies. PH3 levels were significantly higher in the presence of ADNP than its absence (Fig. 9d). Importantly, the PH3 levels could be partially rescued by restoring ADNP in day 6 *Adnp-/-* ESC-derived neurospheres (Fig. 9e).

**Fig. 9.**
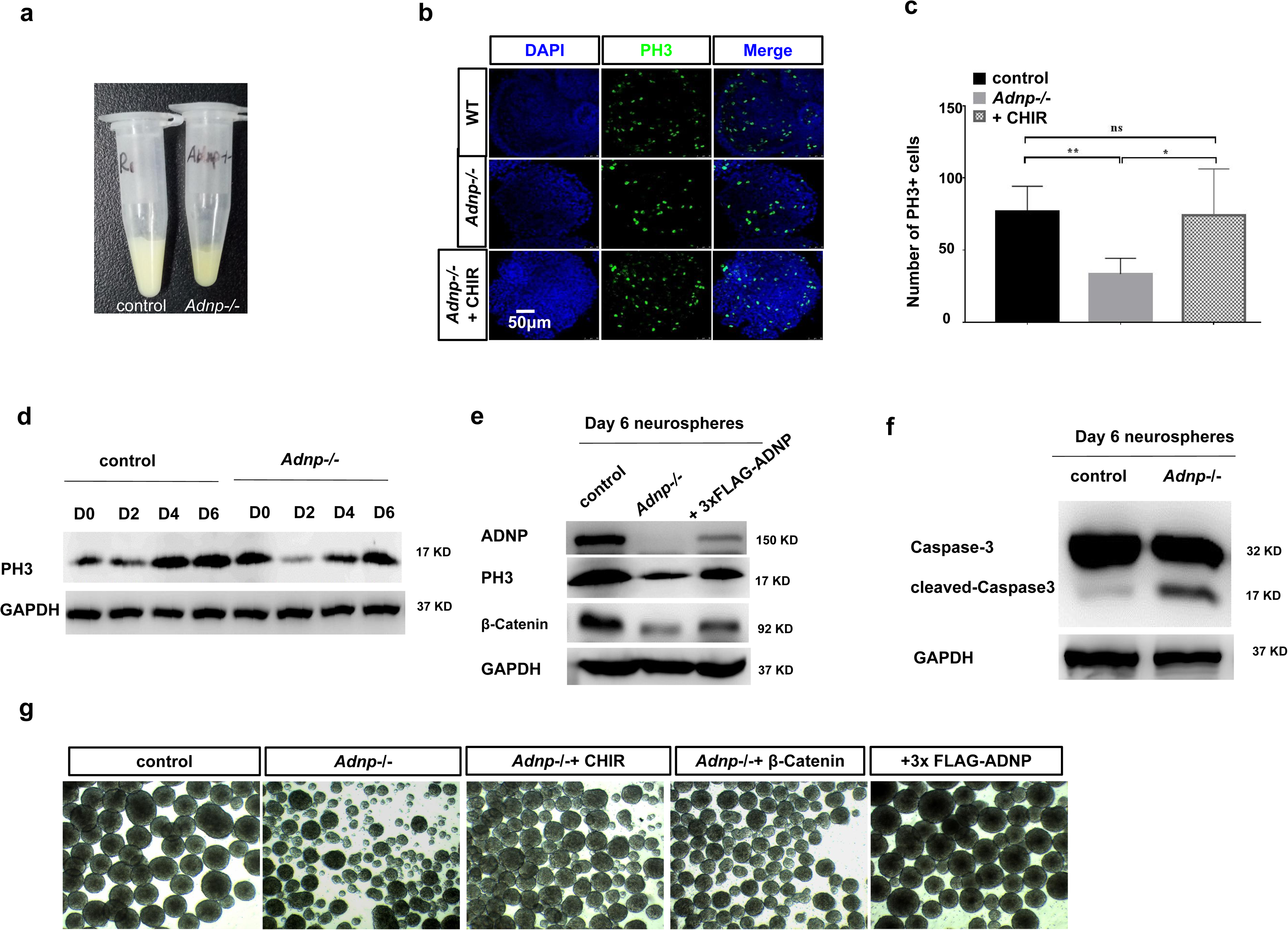
ADNP regulates the proliferation of ESC-derived neural progenitors. **a** Cell mass comparison of day 6 control and *Adnp-/-* ESC-derived neurospheres; **b** Immunostaining of proliferation marker phosphohistone H3 (PH3) for day 6 control and *Adnp-/-* ESC derived neurospheres; **c** Quantification of PH3 positive cells based on panel (**b)**. **d** Representative WB data showing PH3 levels during early stage of control and *Adnp-/-* ESC neural differentiation. **e** Representative WB data showing PH3 levels for day 6 control and *Adnp-/-* ESC-derived neurospheres; **f** WB assay of Caspase-3 for day 6 control and *Adnp-/-* ESCs-derived neurospheres. **g** The morphology of day 6 neurospheres under the indicated conditions (rescue conditions).

To understand whether ADNP was involved in apoptosis during ES cell differentiation towards to a neural fate, WB analysis was performed using Caspase-3 antibodies. Caspase-3 is a well-known key protease that is activated during the early stages of apoptosis. As shown in Fig. 9f, there was an increased cleaved Caspase-3 levels in day 6 *Adnp-/-* ESC-derived neurospheres compared with control ESC-derived neurospheres. This data indicated that loss of ADNP led to the increased apoptosis during ESC neural induction.

Importantly, addition of Wnt signaling agonist CHIR or restoring FLAG-ADNP could partially rescue the cell proliferation defects by loss of ADNP, which supported that ADNP controls cell proliferation or apoptosis by modulating Wnt/β-catenin signaling pathway (Fig. 9b-g). It is well-known that Wnt signaling plays key roles in cell fate determination, differentiation, proliferation and apoptosis^18^.

### Loss of *adnp* leads to the reduced Wnt signaling and defective brain development in a zebrafish model

Zebrafish has been proven to be an excellent in vivo model to study vertebrate neurogenesis and pathology of ASD-related genes^28, 29^. We decided to use zebrafish as an in vivo model to further study the function of ADNP in neural induction and early neurogenesis. Zebrafish has two *adnp* homologues: *adnpa* and *adnpb* gene. Both genes were maternally deposited into the zygote and expressed ubiquitously until the early gastrulation stage when the expression of *adnpa* persisted while the expression of *adnpb* began to disappear. During early somitogenesis, the expression of both genes re-appeared along the dorsal midline, and by 24 hpf was gradually refined to head and gut regions (Supplementary Fig. 8a)^30^. After 72h, *adnpa* expression restricted to the forebrain, the midbrain, the mid-hindbrain boundary and eye area (Supplementary Fig. 8b).

To understand the function of Adnp in zebrafish, we generated *adnpa* and *adnpb* mutants using the CRISPR/Cas9 technology (Fig. 10a, b). Both *adnpa* and *adnpb* mutant embryos can grow up to adults without apparent morphological defects (Supplementary Fig. 8c). We went on to obtain *adnpa adnpb* double mutant zebrafish lines by crossing *adnpa-/-* and *adnpb-/-* fish. We observed that a small portion of *adnpa adnpb* double mutant embryos (about 1/7) at 1 dpf showed abnormal morphology, ranging from slightly ventralization to severe ventralized phenotypes (Supplementary Fig. 8d). The majority of *adnpa adnpb* double mutant embryos appeared normal at 1 dpf (Supplementary Fig. 8d). Strikingly, about 16% (30/190) of these embryos at 48 hpf displayed abnormal morphology with massive neuronal death in brain (Fig. 10c). To further study the neuronal death phenotype, we performed immunofluorescence (IF) staining of HuC/D antibodies for 36 hpf control embryos and mutant embryos that began to show neuronal death in the embryonic brain. As shown in Fig. 10d, the HuC/D signal in *adnpa adnpb* double mutant embryos was greatly decreased compared to that of controls. To further investigate the effects of Adnp deficiency on embryonic brain neuron development, we crossed *adnpa adnpb* double mutant females to the transgenic Tg(*elavl3*: GFP) male fish. We found that the GFP signal was significantly reduced in 3 dpf Adnp deficient embryos compared to the controls (Fig. 10e). Finally, Whole mount in situ hybridization was performed for 2 dpf control and *adnpa adnpb* double mutant embryos to examine the expression of neural developmental genes. We found that the expression of *dlx5a, neurod1* and *phox2a* was greatly reduced in the double mutant embryos compared with sibling controls (Fig. 10f).

**Fig. 10.**
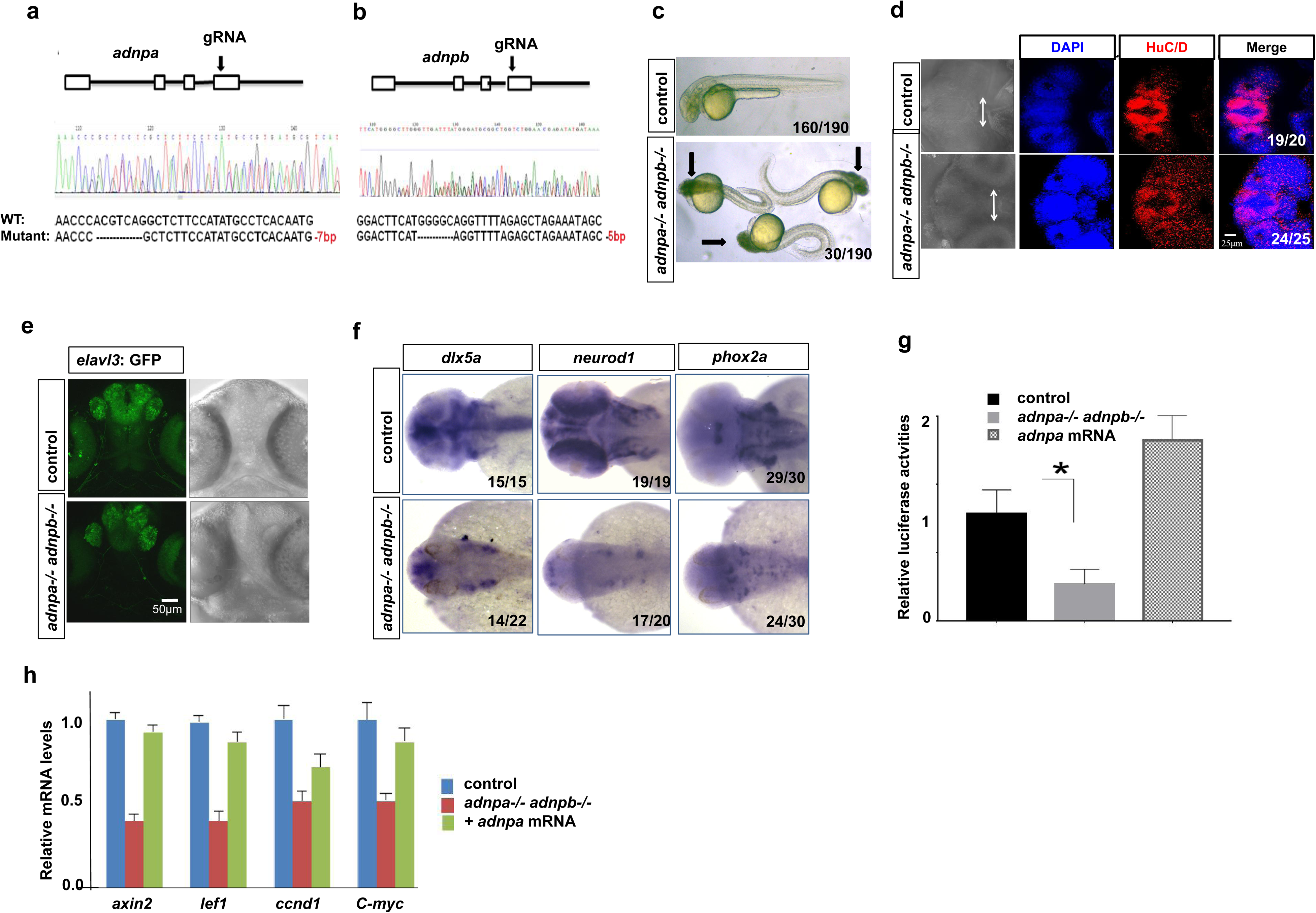
Loss of *adnp* leads to reduced Wnt signaling and defective neural development in zebrafish embryos. **a** CRISPR gRNA design and genotyping of *adnpa* mutant; **b** gRNA design and genotyping of *adnpb* mutant; **c** Representative morphology of 2 dpf control and *adnpa-/- adnpa-/-* double mutants. Black arrow showing the obvious cell death (black area) in head region of mutant embryos. **d** 3D Z-stack image showing the reduced expression of HuC/D in 2 dpf control and double mutant zebrafish brain. Arrow show the distance between the convex tip of the eye cups. **e** GFP fluorescence signal showing the reduced *elavl3*-GFP signal in *adnpa adnpb* deficient embryos. **f** In situ hybridization images for the indicated neural markers for 2 dpf control and double mutant embryos. **g** TopFlash luciferase activity assay for total lysates made from 8 hpf control, double mutant and *adnpa* mRNA overexpressing embryos. **h** The expression of Wnt target genes in 8 hpf control, double mutant and *adnpa* mRNA injected mutant embryos.

Next, we asked whether loss of Adnp in zebrafish embryos led to the reduction of Wnt/β-catenin signaling. We performed TopFlash luciferase assay for lysates from 8 hpf wild type and *adnpa adnpb* double mutant embryos. As shown in Fig. 10g, the TopFlash luciferase activities were greatly reduced in the absence of Adnp. To confirm that Wnt signaling was reduced in *adnpa adnpb* double mutant zebrafish embryos, we examined the expression of known Wnt target genes such as *lef1*, *ccnd1* and *axin2*. qRT-PCR assay showed that the expression of these genes was significantly reduced in *adnpa adnpb* double mutant embryos, which could be partially rescued by the micro-injection of *adnpa* mRNAs (Fig. 10h).

On the other hand, we overexpressed Adnpa by injecting *adnpa* mRNAs into 1-cell stage wild type zebrafish embryos. Overexpression of Adnpa leads to dorsalization of embryos (Supplementary Fig. 10e) and activation of the Wnt/β-catenin signaling pathway as revealed by the TopFlash luciferase reporter assay (Fig. 10g).

Taken together, we concluded that loss of Adnp led to the reduction of Wnt signaling and defective brain development in zebrafish embryos.

## Discussion

De novo mutations in ADNP have recently linked to the Helsmoortel-Van der Aa syndrome, a complex neural developmental disorder displaying multiple symptoms usually manifest in early childhood. In fact, *ADNP* has been proposed as one of the most frequent ASD-associated genes as it is mutated in at least 0.17% of ASD cases. It is therefore important to determine the molecular function of this protein, especially during embryonic neurogenesis.

We found that loss of ADNP had little effect on the mRNA levels of *Ctnnb1* gene which encodes β-Catenin in ESCs. This ruled out that ADNP controls β-Catenin levels by transcriptionally regulating *Ctnnb1*. In fact, *Ctnnb1* gene was barely bound by ADNP as revealed by the ChIP-seq assay (our unpublished observation)^4^. During ESC differentiation towards a neural fate, ADNP translocates from predominantly in the nuclei in ESCs to predominantly in the cytoplasm in neural cell types. These observations suggested that the intra-cellular localization of ADNP is tightly regulated during neural induction, or that ADNP plays an important role in the cytoplasm for the induction, maintenance and mature of neural cell types. By IP in combination with mass spectrometry assay, β-Catenin was highly presented as a top ADNP interacting protein. Considering that Wnt/catenin signaling pathway is critically required for neural induction and differentiation, we reasoned that ADNP may control neural induction or embryonic neurogeneis by regulating Wnt signaling via targeting its key transducer β-Catenin. We speculated that the most likely mechanism was that ADNP controls the stability of β-Catenin in the cytoplasm. This idea was in line with the observation that although ADNP interacts with β-Catenin in both the nuclei and the cytoplasm, loss of ADNP had a significant reduction of the cytoplasmic but not the nuclear faction of β-Catenin. In this work, we provided solid evidences that ADNP directly associates with β-Catenin and regulates its phosphorylation by GSK3β. Mechanistically, ADNP functions to maintain the proper levels of β-Catenin through binding to its armadillo domain which prevents its association with key components of the degradation complex: Axin and APC.

NAP, an 8 aa peptide derived from ADNP, was shown to interact with tubulin to enhance microtubule assembly and neurite outgrowth, and protect neurons and glia exposed to toxins in mouse and rat models^1, 15, 16^. Recently, it was shown that major symptoms of the Helsmoortel-vanderAa syndrome, such as developmental delays, vocalization impediments, motor and pace dysfunctions, and social and object memory impairments, could be efficiently ameliorated by daily NAP administration^17^. Towards the clinical development of NAP (CP201) for the treatment of the Helsmoortel-Van der Aa syndrome, it is important to comprehensively understand the mechanisms underlying the neuroprotective role of NAP. In this work, we showed that ADNP-NAP alone can interact with β-Catenin, suggesting that NAP can execute similar function as ADNP in the regulation of Wnt/β-Catenin signaling pathway. This result provides another possible mechanistic role of NAP in the improvement of symptoms of the Helsmoortel-Van der Aa syndrome: through maintaining Wnt signaling via stabilizing β-Catenin. In the future, it will be interesting to look into this possibility.

*Adnp-/-* mice are early embryonic lethal, displaying severe defects in neural tube closure and brain formation^5^. And E8.5 *Adnp* mutant embryos exhibited reduced expression of *Pax6*. These observations strongly suggested that ADNP plays a key role in embryonic neurogenesis. In this work, using a ESC in vitro neural differentiation system, we showed that ADNP performs an important role at the early stage of neural differentiation by promoting the expression of key neuroectoderm developmental genes. Restoring ADNP levels or activation of Wnt signaling by the small molecule CHIR at the early stage of ESC neural induction can partially rescue the gene expression and morphological defects of day 6 *Adnp-/-* ESC-derived neurospheres, suggesting that ADNP controls the expression of early neural developmental genes through Wnt/β-catenin signaling pathway. In addition, we show that ADNP is abundantly expressed in the whole brain of 10-week-old mice, and is required for proper cerebellar neuron formation. Together, our work suggested that ADNP is cell autonomously required for not only embryonic neurogenesis but also the formation and maturation of vertebrate brain.

The majority of 2dpf *adnpa-/- adnpb-/-* zebrafish embryos exhibited mild phenotype in the brain development. Strikingly, a small portion of the double mutant embryos exhibited massive neuronal cell death in the head region, which recapitulates the key phenotype of *Adnp* deficient mouse embryos. We also showed that ADNP plays important role in cerebellar neuron formation using ESC-based in vitro cerebellar neuron differentiation model. Thus, our work in both ESC and zebrafish models strongly supported the hypothesis that ADNP is required for embryonic brain formation and its loss causes severe neural developmental defects. In the future, more work is needed to test this hypothesis in vivo. Because *Adnp* mutant mice are embryonic lethal, it will be necessary to establish a conditional knockout mouse model to dissect out the function of ADNP in brain formation.

While no single gene is mutated in more than 1% of patients, autism-related genes may functionally converge to commonly affected cellular pathways and protein-protein interaction networks^7^. It is proposed that the most commonly affected networks are the Wnt signaling pathway, and the pathways involving neurogenesis, synaptic function and the chromatin remodeling. An increasing number of studies have shown that ADNP functions as a key chromatin regulator by interacting with chromatin remodelers or regulators. For instances, ADNP interacts with core sub-units of SWI/SNF chromatin remodeling complex such as BRG1 and BAF250 as well as CHD4, a key component of the NuRD complex^4, 7^. Because ADNP is closely involved in the regulation of both the chromatin remodeling and the Wnt signaling pathways, we propose that ADNP is a master regulator implicated in ASD and the related disorders. This may imply why *ADNP* is one of the most frequently mutated genes associated with neural developmental disorders, including the Helsmoortel-Van der Aa syndrome.

## Methods

### ES Cell Culture

Mouse embryonic stem cells (mESCs) R1 were maintained in Dulbecco’s Modified Eagle Medium (DMEM, BI, 01-052-1ACS) high glucose media containing 10% fetal bovine serum (FBS, Gibco, 10099141), 10% knockout serum replacement (KSR, Gibco, 10828028), 1 mM sodium pyruvate (Sigma, S8636), 2 mM L-Glutamine (Sigma, G7513), 1,000 U/ml leukemia inhibitory factor (LIF, Millipore, ESG1107,) and penicillin/streptomycin (Gibco, 15140-122) at 37°C with 5% CO2.

The 2i culture condition was used as described previously^31^. The commercial ESGRO-2i Medium (Merck-Millipore, SF-016-200) was also used when necessary. We found that in 2i medium, *Adnp-/-* ESCs adopted morphology indistinguishable to that of control ESCs, and maintain self-renewal capacity for more than 20 passages that we tested.

### Embryoid body formation

ESCs differentiation into embryoid body was performed in attachment or suspension culture in medium lacking LIF or knockout serum replacement (KSR), as described previously^32^.

### ESC differentiation towards a neural fate

To induce ESC directional neural differentiation in suspension, a neurosphere suspension culture system was used. For Stage 1 of neural induction, cells were dissociated and suspended at a density of 1.5×10^6^ cells in 10 cm^2^ dish coated with 5 ml 1% agarose. The dissociated cells were culture with 45% Neurobasal medium (Life Technologies, 21103-049) and 45% DMEM/F12 (BI, 01-172-1ACS) containing 1×B27 (Gibco, 17504044), 1× N2 (Gibco, 17502048) and 20 ng/ml bFGF (Peprotech, AF-100-18B) and 10 ng/ml EGF (Peprotech, 315-09-1000). Cells were allowed to be cultured on a shaker with low speed at 37 °C in 5% CO2 incubator. Half of the culture medium was changed every two days. For Stage 2 of neuronal differentiation, the neurospheres were plated onto gelatin-coated 6-well plates at day 9, and were cultured for additional 8-10 days in DMEM/F12 containing 2.5% FBS and 1 μM retinoic acid (Sigma, R2625).

ESCs were differentiated into cerebellar neuron cells using an approach adapted from the previously described protocols^24–27^. The differentiation medium (SFEM) was prepared as follows: DMEM supplemented with 5% KSR, 0.1 mM nonessential amino acids (Gibco, 11140050), 1 mM sodium pyruvate, 2 mM L-Glutamine and 0.1 mM 2-mercaptoethanol. For generation of granule and Purkinje progenitor cells, mouse ESCs were dissociated into single cells and suspended at a density of 5.0×10^4^ cells per ml with SFEM. The cells were allowed to aggregate into neurospheres for 5 days. To induce MATH1^+^ cerebellar precursors, final concentration of 6.5 ng/ml BMP4 (Peprotech, 120-05ET-10), 20 ng/ml WNT3a (Peprotech, 315-20-10) and 50 ng/ml Fgf8b (Peprotech, 100-25-25) were added to the medium from day 5-10. For the cerebellar neuron induction, the medium was changed to the Neurobasal medium supplemented with 1×B27 and cells were cultured from day 10 to day 20.

For the inhibition of the proteasome degradation, mESCs or neurospheres were treated with 10 μM MG132 (MCE, HY-13259) for 4 hours before harvesting cells for the next step. For rescue experiments by activating Wnt signaling, mESCs or neurospheres were treated with 25 ng/ml Wnt3a (Peprotech, 315-20-10) or 3 μM CHIR99021 (Selleck, S1263) from the indicated time points during ESC neural induction.

### Lentiviral transduction for shRNA knockdown

The shRNA plasmids for *Adnp* (TRCN0000081670; TRCN0000081671), and the GFP control (RHS4459) were purchased from Dharmacon (USA). To make lentivirus, shRNA plasmids and Trans-lenti shRNA packaging plasmids were co-transfected into H293T cells according to the kit manual (Open Biosystems, TLP4615). After determining the virus titer, mESCs were transduced at a multiplicity of infection of 5:1. Puromycin selection (1 μg/ml) for 4 days was applied to select cells with stable viral integration. Quantitative PCR (qPCR) and Western blot were used to assess the knockdown of *Adnp*.

### Generation of *Adnp-/-* ESCs

*Adnp-/-* mESCs were generated by CRISPR/Cas9 technology. Briefly, we designed two gRNAs on exon 4 of *Adnp* gene by using the online website http://crispr.mit.edu/. The gRNAs sequences are: gRNA 1: 5’-CCCTTCTCTTACGAAAAATCAGG-3’; gRNA 2: 5’-CTACTTGGTGCGCTGGAGTTTGG-3’. gRNAs were cloned into the pUC57-U6 expression vector with G418 resistance. The plasmids encoding gRNA and hCas9 were co-transfected into mESCs using Lipofectamine 2000 (Gibco, 11668019). After 48 hours, mESCs were selected with 500 μg/ml G418 for 7 days. Then the cells were re-seeded on a 10-cm dish coated with 0.1% gelatin to form colonies. The single colony was picked up, trypsinized and passaged at low density. DNA from single colonies from the passaged cells was extracted and used for genotyping.

### Generation of 3×Flag Tagged *Adnp-/-* mESC Cell Lines

The full-length *Adnp* cDNA (NM_009628.3) was amplified by PCR and then cloned into pCMV-3×Flag vector. The full-length *Adnp* cDNA sequence containing N-terminal 3×Flag sequence was subcloned into pCAG-IRES-Puro vector. To make stable transgenic cells, *Adnp-/-* mESCs were transfected with pCAG-IRES-Puro-3×Flag-Adnp vector using Lipofectamine 2000 (Gibco, 11668019). 48 hours later, cells were selected by 1 μg/m l puromycin. After 4-5 days drug selection, cells were expanded and passaged. Western Blot assay was performed to confirm the transgenic cell line using a FLAG antibody.

To make Tet-Express inducible transgenic cells, 3×Flag-*Adnp* or HA-*Ctnnb1* were subcloned into pTRE3G or pLVX-tight-puro expression vector (Clontech, PT5167-1 and PT3996-5) using In-fusion HD cloning system (Clontech, 638910). Stable transgenic cell lines were established according to the manual of the Tet-Express inducible expression systems (Clontech, 631169). Briefly, *Adnp-/-* ESCs were transfected with 2 μg pTRE3G-3×Flag-*Adnp* or pTRE3G-HA-*Ctnnb1* with linear 100 ng puromycin marker using Lipofectamine 2000 transfection reagent. 96h later, 1 μg/ml puromycin was added and drug selection was performed for two weeks to establish the stable transgenic cell line. To induce target gene expression, 3×10^6^ transgenic cells were plated in 6-well plates. The next day, the Tet-Express transactivitor (Clontech, 631178) was added (3 μl Tet-Express to a final 100 μl total volume according to the kit manual) for 1 h in serum-free medium to induce the target gene expression. Then cells were allowed to grow in complete medium for an additional 12–24 h before assay for the target protein induction. Western blot were used to assess the target protein expression levels using a FLAG or HA antibodies. In the absence of Tet-Express transactivitor, pTRE3G provides very low background expression, whereas addition of Tet-Express proteins strongly transactivates target genes.

### FACS experiments

Neurospheres and SFEB cell aggregates were washed with DPBS. The aggregates were dissociated into single cells with 0.25% trypsin (BI, 03-050-1A). After fixing with 4% paraformaldehyde at room temperature for 10 minutes, the cells were washed with 0.5% PBSA (0.5% BSA in PBS) and treated with 90% cold methanol for 15 minutes. After extensive washes, the cells were suspended with 200 μl PBSA and filtered with the flow tube. Then the cells were incubated with anti-NESTIN (Abcam, ab7659), anti-PAX6 (Proteintech, 12323-1-AP) antibody at room temperature for 15 minutes. After three-times wash with 0.5% PBSA, the cells were incubated with secondary antibodies (1: 500 dilution in PBSA, AlexaFluor 488) at room temperature for 15 minutes in the dark. The cells were washed with PBSA for two-times and resuspended with 400 μl PBSA, and were analyzed by the BD AccuriC6 flow cytometer.

### RNA preparation, RT-qPCR and RNA-seq

Total RNA from mESCs, neurospheres and neuronal cells was extracted with a Total RNA kit (Omega, R6834-01). A total of 1 μg RNA was reverse transcribed into cDNA using the TransScript All-in-One First-Strand cDNA synthesis Supermix (Transgen Biotech, China, AT341). Quantitative real-time PCR (RT-qPCR) was performed using the TransStart® Tip Green qPCR SuperMix (Transgen Biotech, China, AQ-141). The primers used for RT-qPCR were listed in Table 1. All experiments were repeated for at least three times. The relative gene expression levels were calculated based on the 2-ΔΔCt method. Data were shown as means ± S.D. The Student’s t test was used for the statistical analysis. The significance is indicated as follows: *, p < 0.05; **, p < 0.01; ***, p < 0.001.

**Table 1.**
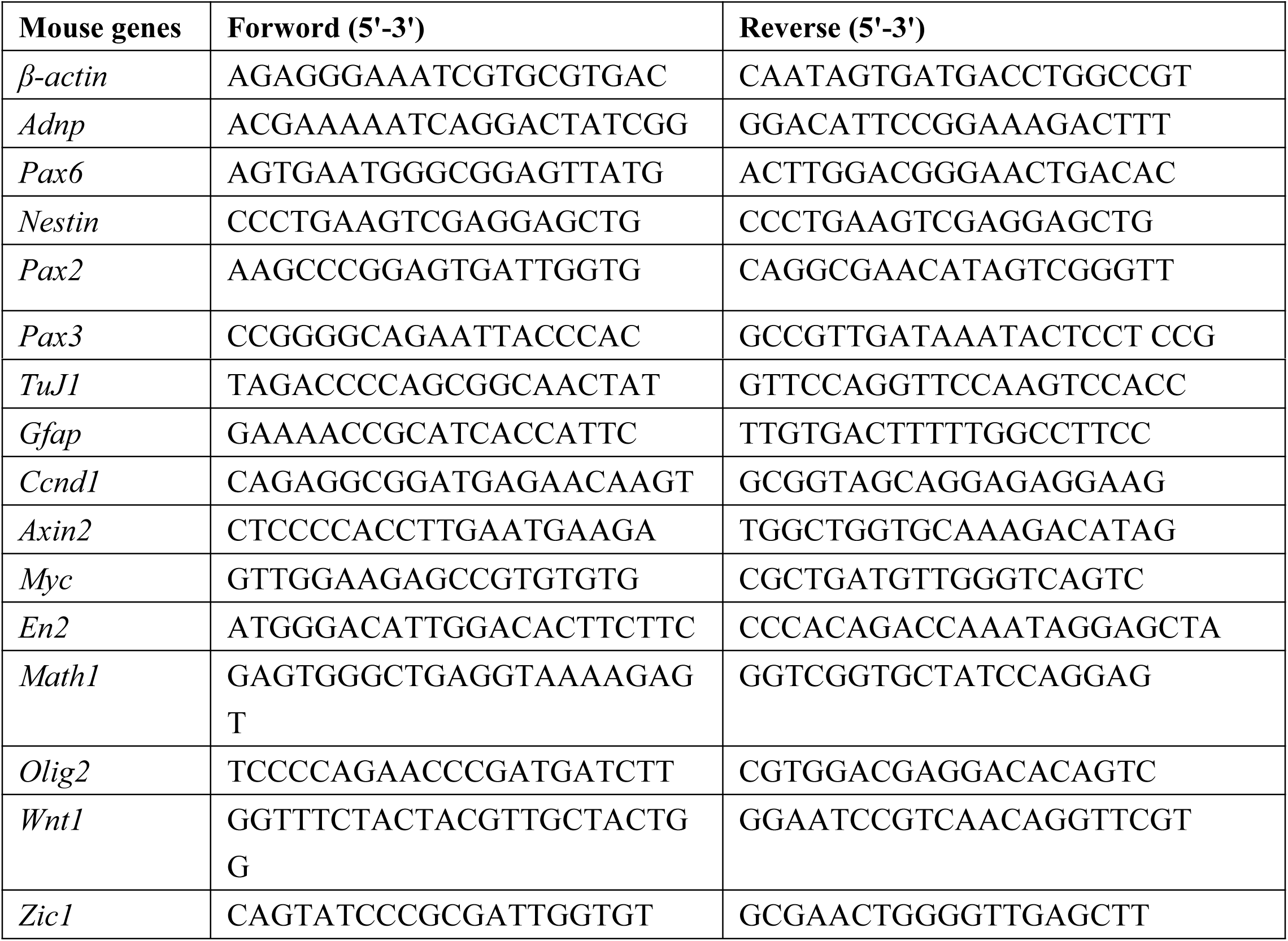
The primers for RT-qPCR analysis.

**Table 2.**
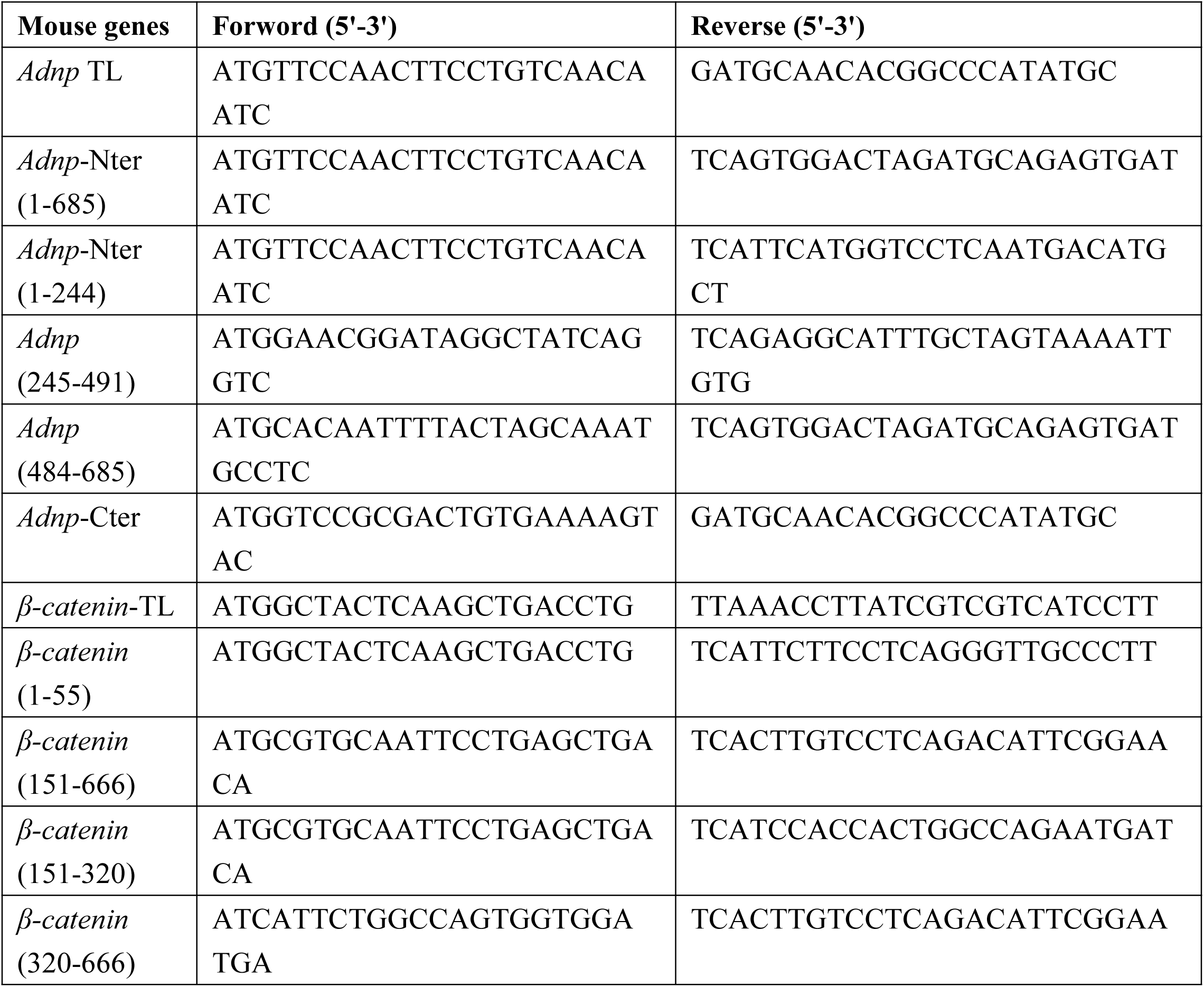
The primers for cDNA cloning.

For RNA-seq, ESCs and day 3-, day 4- and day 6-control and mutant ESC-derived neurospheres were collected and treated with Trizol (Invitrogen). RNAs were quantified by a Nanodrop instrument, and sent to BGI Shenzhen (Wuhan, China) for making RNA-seq libraries and deep sequencing. For each time points, at least two biological repeats were sequenced. DEGs were defined by FDR < 0.05 and a Log2 fold change >1.5 fold was deemed to be differentially expressed genes (DEGs).

### Protein extraction and Western blot analysis

For protein extraction, ESCs cells and neurospheres were harvested and lysed in TEN buffer (50 mM Tris-HCl, 150 mM NaCl, 5 mM EDTA, 1% Triton X-100, 0.5% Na-Deoxycholate, supplement with Roche cOmplete™ Protease Inhibitor). The lysates were quantified by the Bradford method and equal amount of proteins were loaded for Western blot assay. Antibodies used for WB were ADNP (AF5919, R&D Systems), non-phospho (Active) β-Catenin (Cell Signaling Technology CST, #8814), Phospho-β-Catenin (Ser33/37/Thr41) (CST, #9561), total β-Catenin (CST, #9562), anti-NESTIN (Abcam, ab7659), anti-PAX6 (Proteintech, 12323-1-AP), anti-PCNA (Cusabio, CSB-PA01567A0Rb), anti-GSK3β (Proteintech, 2204-1-AP), anti-pGSK3β (CST, 9336S), anti-PH3 (Santa Cruz, sc-8656-R), anti-Caspas3 (BD, 559565), anti-L7 (Takara, M202), anti-PAX2 (Abcam, ab79389), anti-TuJ1 (Abcam, ab78078), anti-GFAP (Abcam, ab53554), anti-HuC/D (Life Technologies, A21271), anti-β-Catenin (Sigma, C7207), Anti-FLAG (F3165, Sigma, 1:1000), anti-MYC antibody (Transgen Biotech, HT101) and anti-HA (Abbkine, A02040, 1:1000). Briefly, the proteins were separated by 10% SDS-PAGE and transferred to a PVDF membrane. After blocking with 5% (w/v) non-fat milk for 1 hour at room temperature, the membrane was incubated overnight at 4°C with the primary antibodies. Then the membranes were incubated with a HRP-conjugated goat anti-rabbit IgG (GtxRb-003-DHRPX, ImmunoReagents, 1:5000), a HRP-linked anti-mouse IgG (7076S, Cell Signaling Technology, 1:5000) for 1 hour at room temperature. The GE ImageQuant LAS4000 mini luminescent image analyzer was used for photographing.

### Co-immunoprecipitation assay (Co-IP)

Co-immunoprecipitations were performed with the Dynabeads Protein G (Life Technologies, 10004D) according to the manufacturer’s instructions^33^. Briefly, 1.5 mg Dynabeads was conjugated with antibodies or IgG overnight at 4 °C. Antibodies were used are: 10 μg IgG (Proteintech, B900610), or 10 μg anti-ADNP antibody (R&D Systems, A5919), or 10 μg anti-FLAG antibody (Proteintech, 20543-1-AP), or 10 μg anti-HA antibody (Abbkine, A02040) or 10 μg anti-MYC antibody (Transgen Biotech, HT101). The next day, total cell lysates and antibody-conjugated Dynabeads were incubated overnight at 4 °C with shaking. After three-times wash with PBS containing 0.1% Tween, the beads were boiled at 95 °C for 5 minutes with the 6×Protein loading buffer and the supernatant was collected for future WB analysis.

### Luciferase reporter assays

TopFlash-luc reporter was kindly provided by Prof. Zongbin Cui from Institute of Hydrobiology. HEK293T cells (about 1×10^5^ cells) were seeded in 24-well plates in DMEM medium containing 10% FBS (Transgen Biotech, FS101-02). After 24 hour, the cells were transiently transfected with the indicated luciferase reporters using Liposomal Transfection Reagent (Yeasen, 40802ES03). For mESCs transfection, the Neon transfection system was used according to the manual. Briefly, the trypsinized mESCs were resuspended in the electroporation buffer at a density of 1×10^7^cells/ml. The TopFlash-luc reporter and pTK-Renilla plasmids were added into cells. Then electroporation was performed by the Neon transfection system according to the manufacture’s instructions. After transfection, the cells were treated with or without 10 ng/ml Wnt3a for 16 hours. The luciferase activity was measured with Dual-luciferase Reporter Assay System (Promega, Madison, E1910). Renilla was used as an internal control.

For luciferase reporter assays in zebrafish embryos, 1-cell stage wild type and *adnpa adnpb* double mutant embryos were injected with 200 pg TopFlash and 20 pg pTK-Renilla plasmids, and the late gastrula stage embryos were collected. Three pools of 10 embryos each were lysed with passive lysis buffer and assayed for luciferase activity according to the Dual luciferase system (Promega). The Topflash luciferase activities were measured with the Dual-luciferase Reporter Assay System. All luciferase reporter assays represent the mean ± standard error from 3 independent measurements of pools. At least three independent experiments were performed in different batches of embryos.

### Immunofluorescence assay

Neurospheres and SFEB aggregates were collected and fixed with 4% paraformaldehyde for half an hour at room temperature. Then the cells were washed with PBST (phosphate-buffered saline, 0.1% Triton X100) for three times, each for 15 minutes. Following the incubation with blocking buffer (5% normal horse serum, 0.1% Triton X-100, in PBS) for 2 hours at room temperature, the cells were incubated with primary antibodies at 4 °C overnight. After three-times wash with PBST, the cells were incubated with secondary antibodies (1: 500 dilution in blocking buffer, Alexa Fluor 488, Life Technologies) at room temperature for 1 hour in the dark. The nuclei were counter-stained with DAPI (Sigma, D9542, 1:1000). After washing with PBS for twice, the slides were mounted with 100% glycerol on histological slides. Images were taken by a Leica SP8 laser scanning confocal microscope (Wetzlar, Germany).

### Immunohistochemistry

Ten-week-old mice brain frozen sections were pretreated with an antigen unmasking solution (10mM Sodium Citrate, pH6.0) for 15min at 95 °C in oven. After that, sections were naturally cooled to room temperature and blocked with 3% normal donkey serum adding 0.1% Tween-20 at room temperature for 2h. The tissue was then incubated overnight at 4 °C with the anti-ADNP antibody diluted in blocking buffer, followed by avidin-biotin-peroxidase complex (1:50; ABC kit, VECTASTAIN Elite ABC system, Vector Labs, Burlingame, CA, USA). Peroxidase was reacted in 0.03% 3,3’-diaminobenzidine (DAB) and 0.003% H_2_O_2_ in PBS. Sections were dehydrated, cleared in xylene, and mounted in neutral balsam. Measuring and quantification of IHC images were performed using the Image-pro Plus 6.0 software (Media Cybernetics).

### Immunoprecipitation in combination with mass spectrometry

For Mass Spectrometry, the IP samples (immunoprecipitated by IgG or ADNP antibody) were run on SDS-PAGE gels and stained with Coomassie Blue. Then the entire lanes for each IP samples were cut off and transferred into a 15 ml tube containing deionized water. The treatment of the samples and the Mass Spectrometry analysis were done by GeneCreate Biological Engineering Company (Wuhan, China).

### Protein-protein interaction assay using a rabbit reticulocyte lysate system

Protein-protein interaction assay using a rabbit reticulocyte lysate system has been described previously^33^. Tagged-ADNP, Tagged-ADNP mutants, Tagged-β-Catenin and Tagged-β-Catenin mutants were synthesized using the TNT coupled reticulocyte lysate system according to the manual (Promega, L5020, USA). Briefly, 1 μg of circular PCS2-version of plasmids were added directly to the TNT lysates and incubated for 1.5 hours at 30 °C. 1 μl of the reaction products were subjected to WB assay to evaluate the synthesized protein. For protein-protein interaction assay, 5-10 μl of the synthesized HA or FLAG tagged proteins were mixed in a 1.5 ml tube loaded with the 300 μl TEN buffer, and the mixture was shaken for 30 minutes at room temperature. Next, IP or pull-down assay was performed using Dynabeads protein G coupled with anti-FLAG or anti-HA antibodies as described above.

### Zebrafifish Maintenance

Zebrafish (*Danio rerio*) were maintained at 28.5 °C on a 12 h light/12 h dark cycle. All procedures were performed with the approval of the Institute of Hydrobiology, Chinese Academy of Sciences, Wuhan, China.

### Generation zebrafish *adnp* mutants by CRISPR/Cas9

Zebrafish *adnp* mutants were generated by CRISPR/Cas9 technology. The gRNA targeting sequences for *adnpa* and *adnpb* gene were 5’- GGACTCTGGAAACCCACGTC -3’ and 5’- GGAGGACTTCATGGGGCAG - 3’, respectively. Guide RNAs (gRNAs) were generated using the MEGAshortscript T7 kit (Thermo Fisher). The Cas9 mRNA was synthesized using the mMESSAGE mMACHINE® SP6 Kit (AM1340, Thermo Fisher). The mixture containing 200 ng/µl Cas9 mRNA and 80 ng/µl guide RNA was co-injected into 1-cell stage zebrafish embryos. The genomic DNA of 20 injected embryos at 24 hpf was extracted and subjected to PCR amplification. The DNA fragments containing the gRNA targeting sequences were amplified by PCR using primers 5’- TGCTCAGATTGCCCGTTT -3’ and 5’-GATAGGTGCACTTCTTGCAGTA -3’ for *adnpa* gene, 5’-AATGTGCACAGCGAGGACTT-3’ and 5’- GACAAGGACTGTGTAGCCCC -3’ for *adnpb* gene, respectively. The genotype was confirmed by DNA sequencing. Adults raised up from injected embryos were screened for mosaic founders by the amplicon sequencing. The mosaic founders were outcrossed to wild type to obtain the F1 offsprings with stable germline transmission. The F1 heterozygous carrying 7 bp deletion (*adnpa*) and 5 bp deletion (*adnpb*) were outcrossed to wild type to generate F2 heterozygous, respectively. The F2 heterozygous were inter-crossed to generate homozygous *adnpa*^-/-^ and *adnpb*^-/-^, respectively. The *adnpa*^-/-^ and *adnpb*^-/-^ adults were outcrossed to each other to generate *adnpa* and *adnpb* double mutant. The subsequent screening was performed by high resolution melt analysis (HRMA) using primers 5’-TTCCGCAATGTTCACAGGGA-3’ and 5’- GGCATATGGAAGAGCCTGACG-3’ for *adnpa*, 5’- CACATCAGGTTGTTTCATATGCCT -3’ and 5’- TCACGTGTCTTTTCCAGACGA -3’ for *adnpb*, respectively.

### Microinjections into zebrafish embryos

For overexpression study, full-length cDNA encoding Adnpa (NC_007122.7) was amplified and cloned into the PCS2 (+) vector. The primers we used are as followed GATGTTTCAGCTTCCAGTGAATAACC (Forward), and CACAAGCCATCATCTACCAAGC (Reverse). The *Adnpa* mRNA was transcribed with mMESSAGE mMACHINE SP6 Kit (Invitrogen, AM1340) and was dissolved in nuclease-free water to the final concentration of 150 ng/μl. A total of 1-5 nl mRNAs were microinjected into one-cell stage wild-type zebrafish embryo using the WPI microinjector (WPI, USA). The 1-2 dpf embryos were anesthetized in 0.016% tricaine and photographed under a Leica stereomicroscope.

### Whole-mount immunofluorescence

The whole-mount immunofluorescence experiments were performed as described^33^. Briefly, 48 hpf zebrafish embryos were fixed in 4% paraformaldehyde overnight at 4 °C and then underwent proteinase K treatment for 8 minutes at RT. The primary antibody we used was anti-HuC/HuD monoclonal antibody (Life Technologies, USA, 1: 350 dilution in PBST). After blocking for 2 h at room temperature with solution (0.1% Triton X-100, 1% BSA, and 1% DMSO in PBS), the embryos were incubated overnight with the primary antibodies at 4 °C with shaking. After three-times wash with PBST, embryos were incubated with the secondary antibodies (1:1000, Goat anti-Mouse IgG secondary antibody, Alexa Fluor 488, Life technologies, USA) for 2 h at RT. Finally, the embryos were counter-stained with DAPI. Embryos were imaged in 4% methyl-cellulose using a Leica SP8 Confocal microscope.

### Whole-mount in situ hybridization (WISH)

WISH was performed as described previously^33^. For making the RNA probes, total RNA was extracted from zebrafish embryos, and cDNAs were made by RT-PCR. Templates were amplified by PCR using primers with SP6 or T7 promoters. Primers we used are listed in Table 3. The DIG-labeled probes for *adnpa*, *adnpb*, *dlx5a*, *neurod1* and *phox2a* were made using the DIG RNA Labeling Kit (SP6/T7) (Roche). Zebrafish embryos at different developmental stages were collected and fixed with 4% PFA overnight at 4 °C. Following the WISH, the embryos were transferred to 6-well plates and submerged in 100% glycerol for imaging.

**Table 3.**
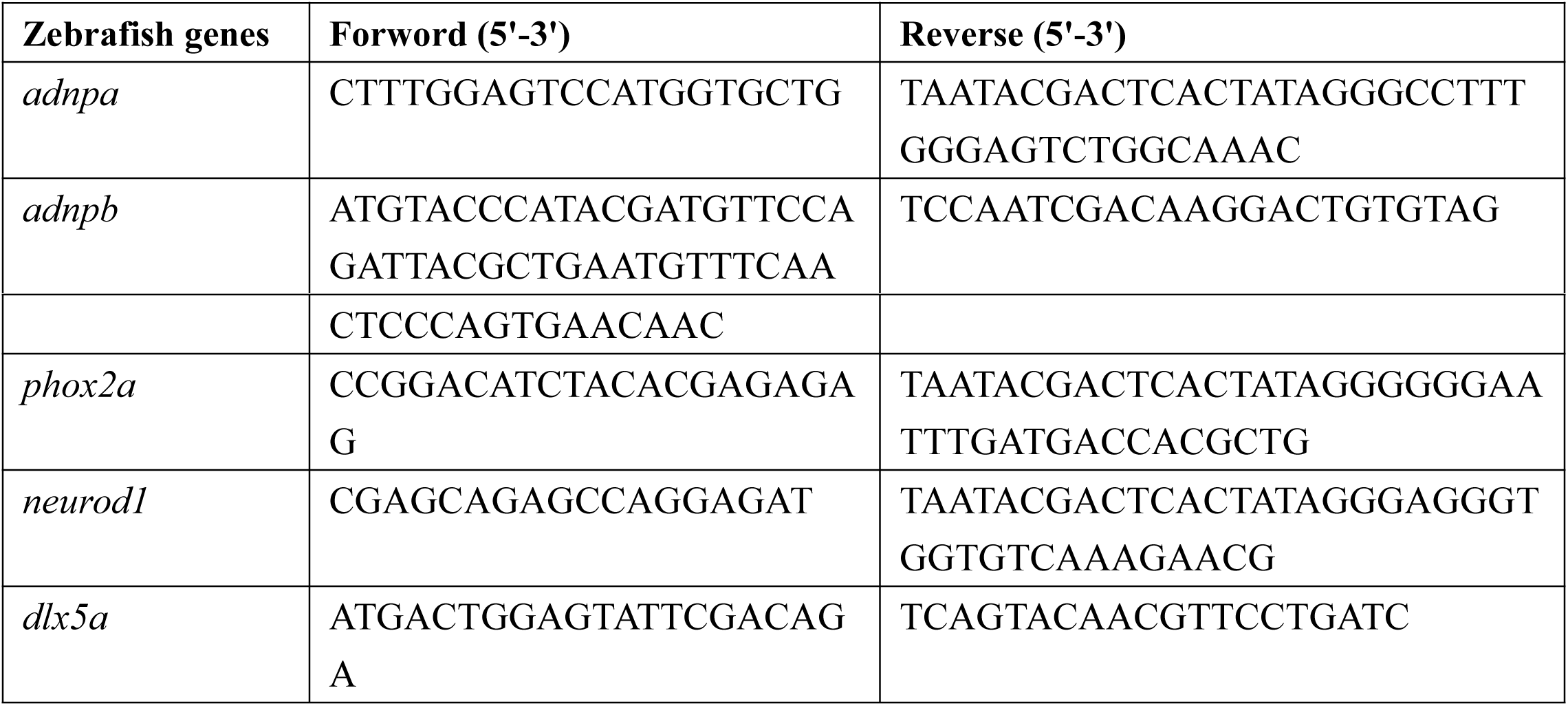
The primers for making In situ hybridization probes.

### Quantification and statistical analysis

Data are presented as mean values ±SD unless otherwise stated. Data were analyzed using Student’s t-test analysis. Error bars represent s.e.m. Differences in means were statistically significance when p< 0.05. Significance levels are: *p< 0.05; **P<0.01.

### Data availability

All data will be available upon request.

## Acknowledgements

This work was supported by National Key Research and Development Program (2016YFA0101100), National Natural Science Foundation of China (31671526), and Hundred-Talent Program (CAS and IHB) (Y623041501) to YH Sun.

## Author contributions

XY Sun performed the stem cell part of work; XX Peng and YQ Cao performed the zebrafish part of work; Y Zhou and Yuhua Sun designed the work; Yuhua Sun provided the final support of the work, and wrote the paper.

---

**Supplementary Fig. 1.**
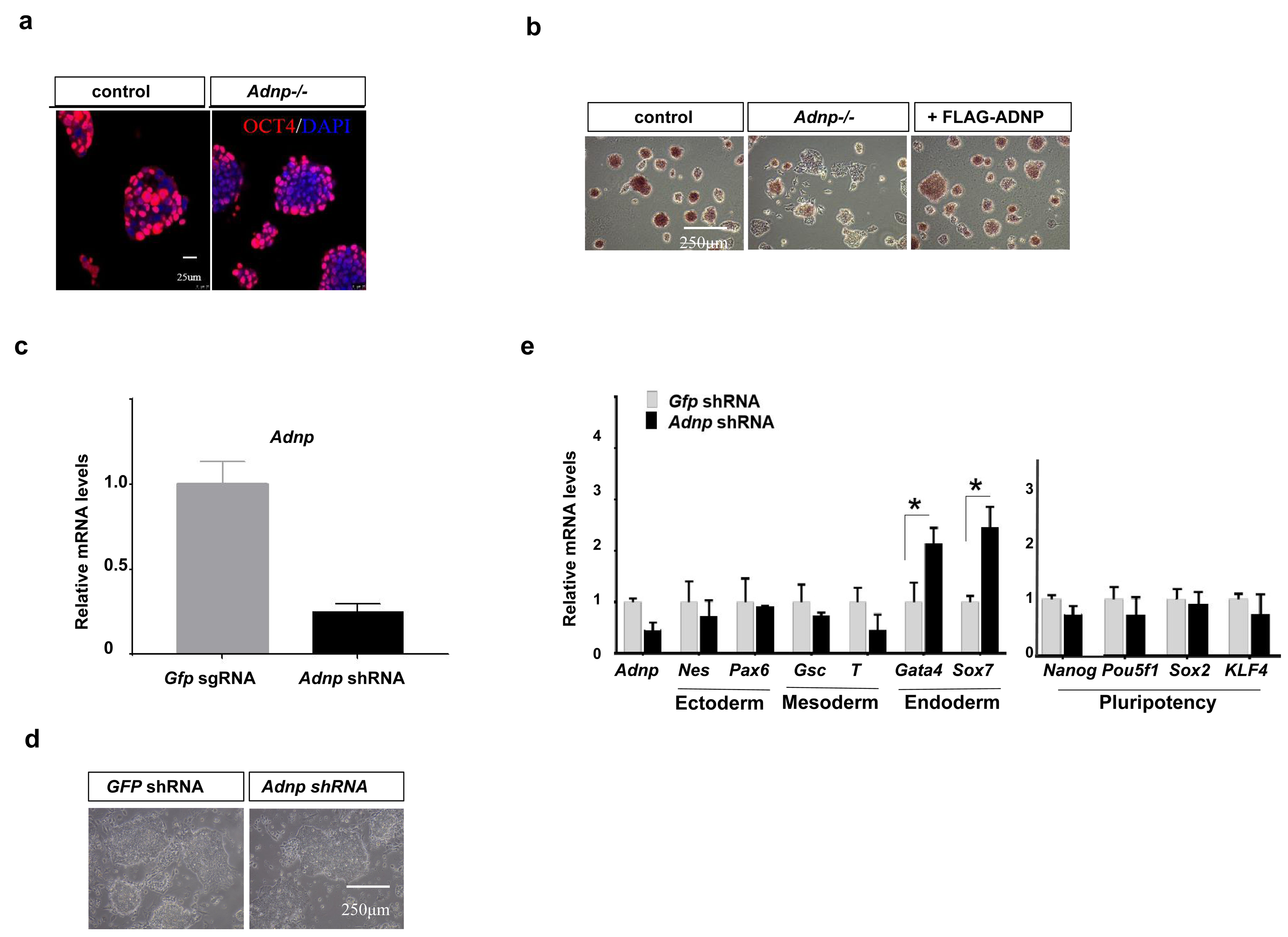
Related to Figure 1. **a** IF staining of OCT4 for control and *Adnp-/-* ESCs. **b** Alkaline phosphotase staining for long-term passaged control and *Adnp-/-* ESCs (shown is successfully passaged for 10 times in LIF/KSR medium). **c** Relative *Adnp* mRNA levels for *Gfp* shRNA and *Adnp* shRNA knockown ESCs. **d** Representative image showing morphology of control and early passaged *Adnp* shRNA ESCs. **e** The expression of representative pluripotency-related, mesodermal, neuroectodermal, endodermal genes in control and early passaged *Adnp* shRNA ESCs. qPCR has been repeated for at least three times and represented as mean values± s.d.

**Supplementary Fig. 2.**
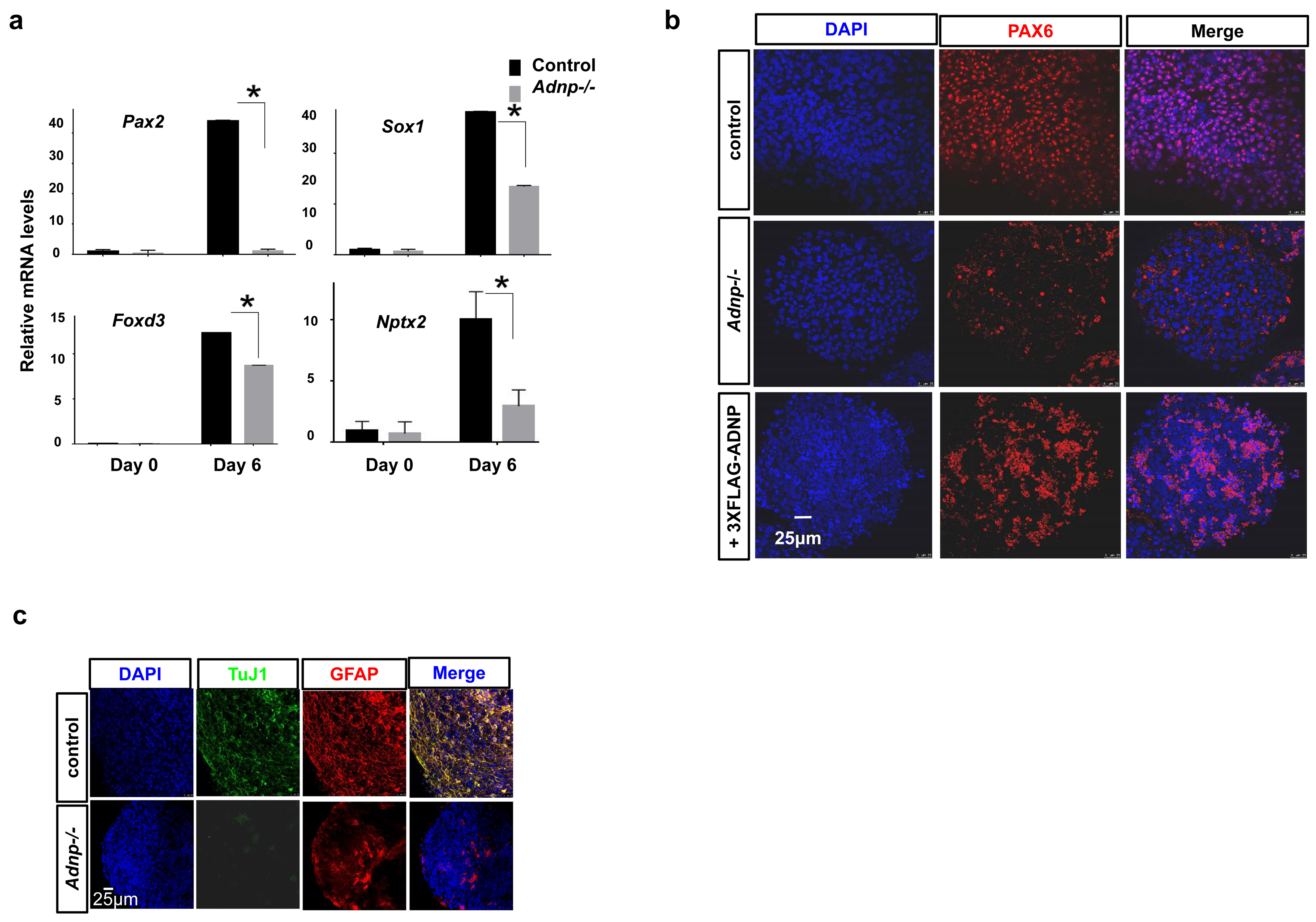
Related to Figure 2. **a** The expression of representative neural genes at indicated time points. **b** IF staining of PAX6 for day 6 control, *Adnp-/-* and FLAG-ADNP restoring *Adnp-/-* ESC-derived neurospheres. **c** IF staining of TuJ1 and GFAP for day 16 control and *Adnp-/-* ESC-derived neuronal cell cultures.

**Supplementary Fig. 3.**
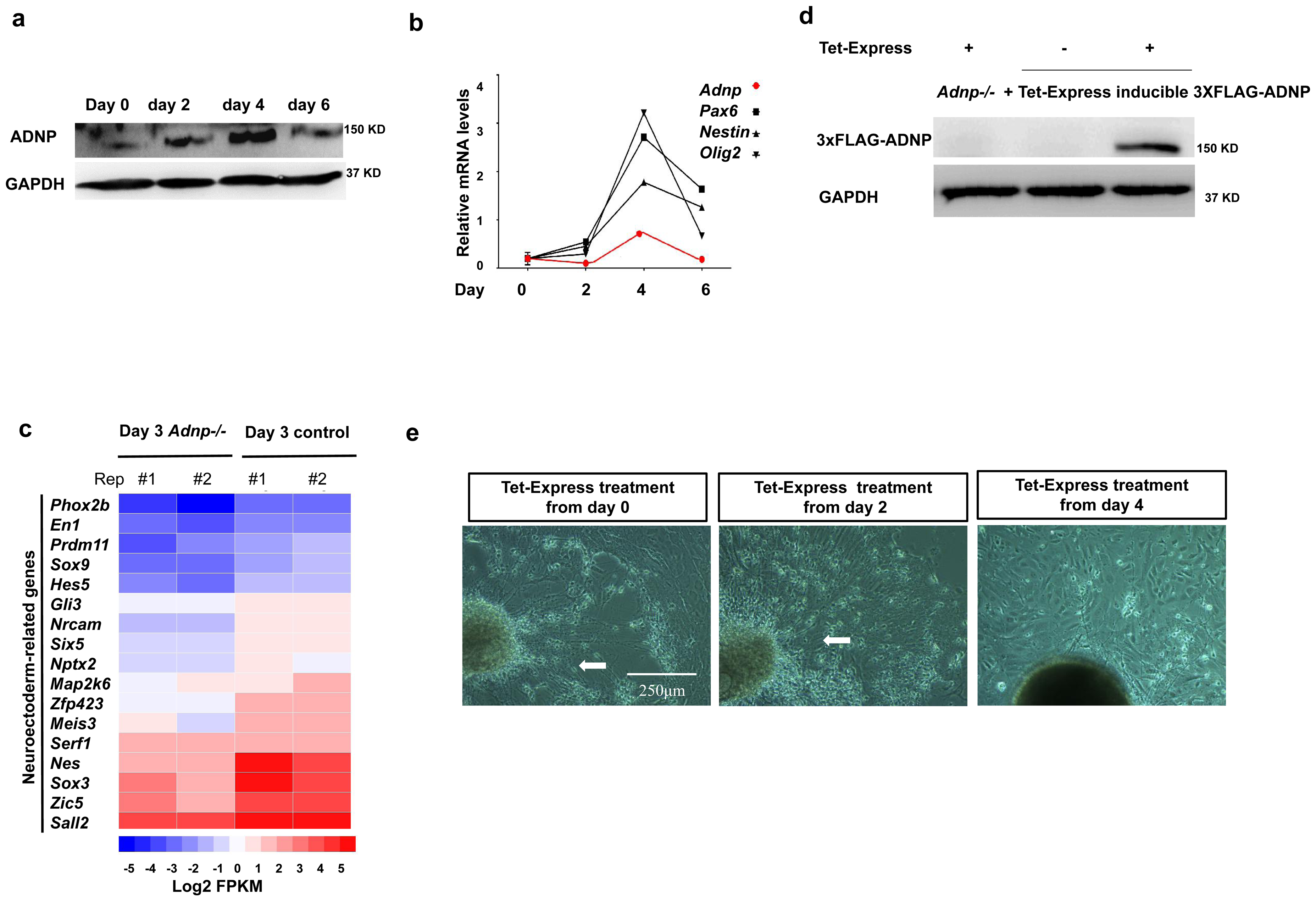
Related to Figure 3. **a** Dynamic expression profile of ADNP during first 6 days of neural induction of ESCs. Experiments were repeated for two times. **b** Dynamic expression profile of *Adnp*, *Pax6*, *Nestin* and *Olig2* mRNAs during the first 6 days of neural induction of wild type ESCs. Experiments were repeated for three times. **c** Heatmap illustrating the expression of selected neuroectoderm genes that were shown as log2 FPKM in day 3 control and *Adnp-/-* ESC-derived neurospheres. Each lane corresponds to an independent biological RNA-seq sample. **d** WB showing that addition of Tet-Express transactivated 3×FLAG-ADNP in *Adnp-/-* ESCs. **e** Representative morphology showing the neuronal fibre structure in *Adnp-/-* ESC-derived day 19 neuronal cell types that added with Tet-Express from day 0, day 2 and day 4 of *Adnp-/-* ESC neural induction. Experiments were repeated for three times, and shown are the representative images.

**Supplementary Fig. 4.**
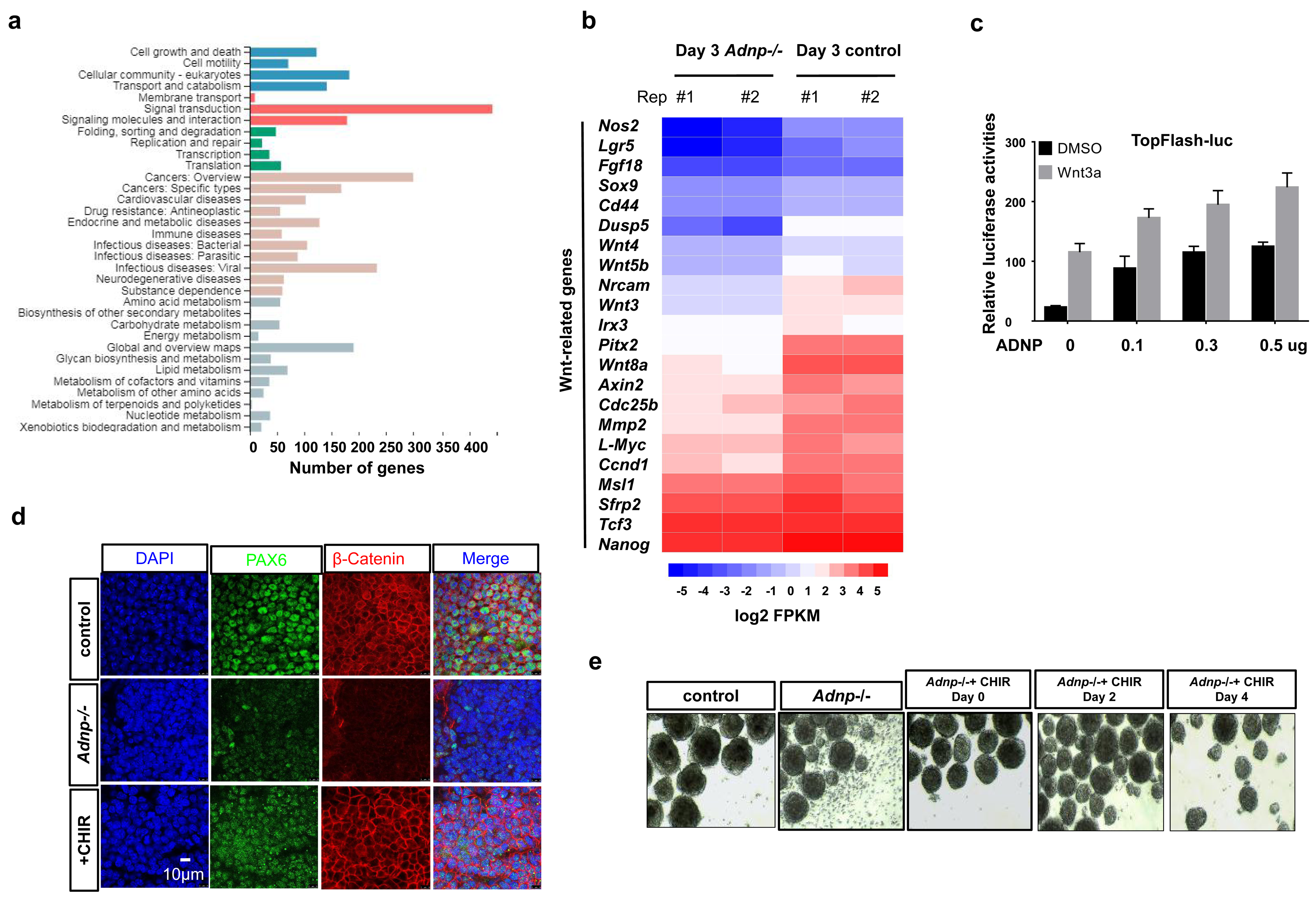
Related to Figure 4. **a** KEGG analysis of of DEGs from day 3 and day 6 control and *Adnp-/-* ESC-derived neurospheres, showing enrichment of pathway related to signaling transduction. **b** Heatmap illustrating the expression of selected Wnt-related genes that were shown as log2 FPKM in day 3 control and *Adnp-/-* ESC-derived neurospheres. Each lane corresponds to an independent biological RNA-seq sample. **c** TopFlash luciferase activity analysis of 293T cells transfected with increasing dose of plasmids encoding ADNP in the absence or presence of Wnt3a. Experiments were repeated for two times. **d** Rescue of PAX6 expression by addition of CHIR. Representative IF staining showing PAX6 signal for day 6 control and *Adnp-/-* ESC-derived neurospheres. **e** Representative morphology of day 6 *Adnp-/-* ESC-derived neurospheres after addition of CHIR from the indicated time points. Experiments were repeated for two times.

**Supplementary Fig. 5.**
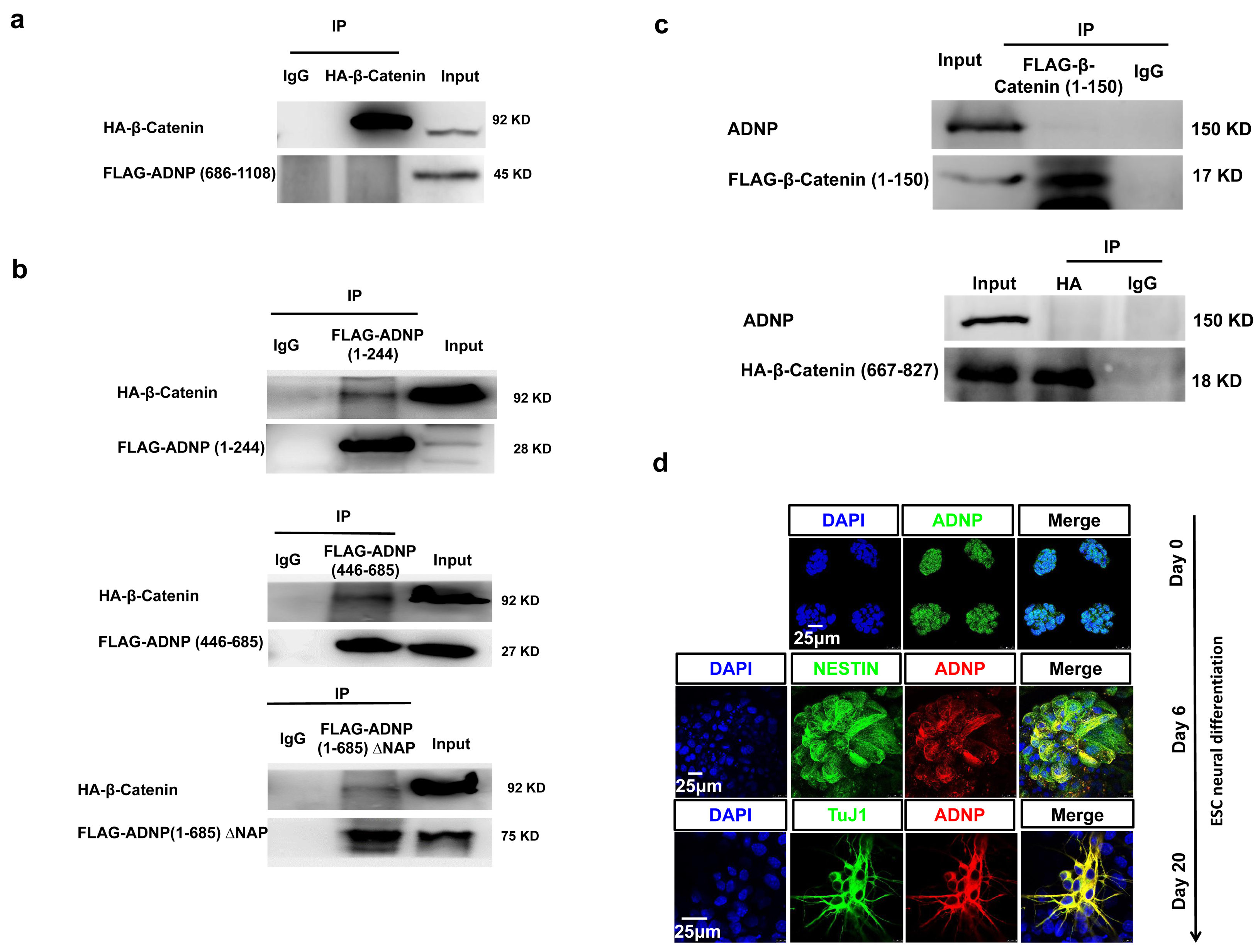
Related to Figure 5. **a** WB showing that the C-terminal of ADNP (686-1108) does not interact with β-Catenin in 293T cells. **b** Dissection of FLAG-ADNP-Nter (1-685) that is responsible to interact with β-Catenin in 293T cells. ADNP fragment 1-224, 446-685 both can interact with β-Catenin. And ADNP-Nter (1-685) without NAP can still interact with β-Catenin. **c** WB showing that β-Catenin (1-150) barely interacts with ADNP in 293T cells. **d** IF immunostaining of ADNP showing that during ESC neural differentiation ADNP translocates from nuclei to cytoplasm, and that ADNP was colocalized with NESTIN and TuJ1 in day 6 ESC-derived neurospheres and day 19 neuronal cell types, respectively. All WB and IF staining experiments have been repeated for at least two times.

**Supplementary Fig. 6.**
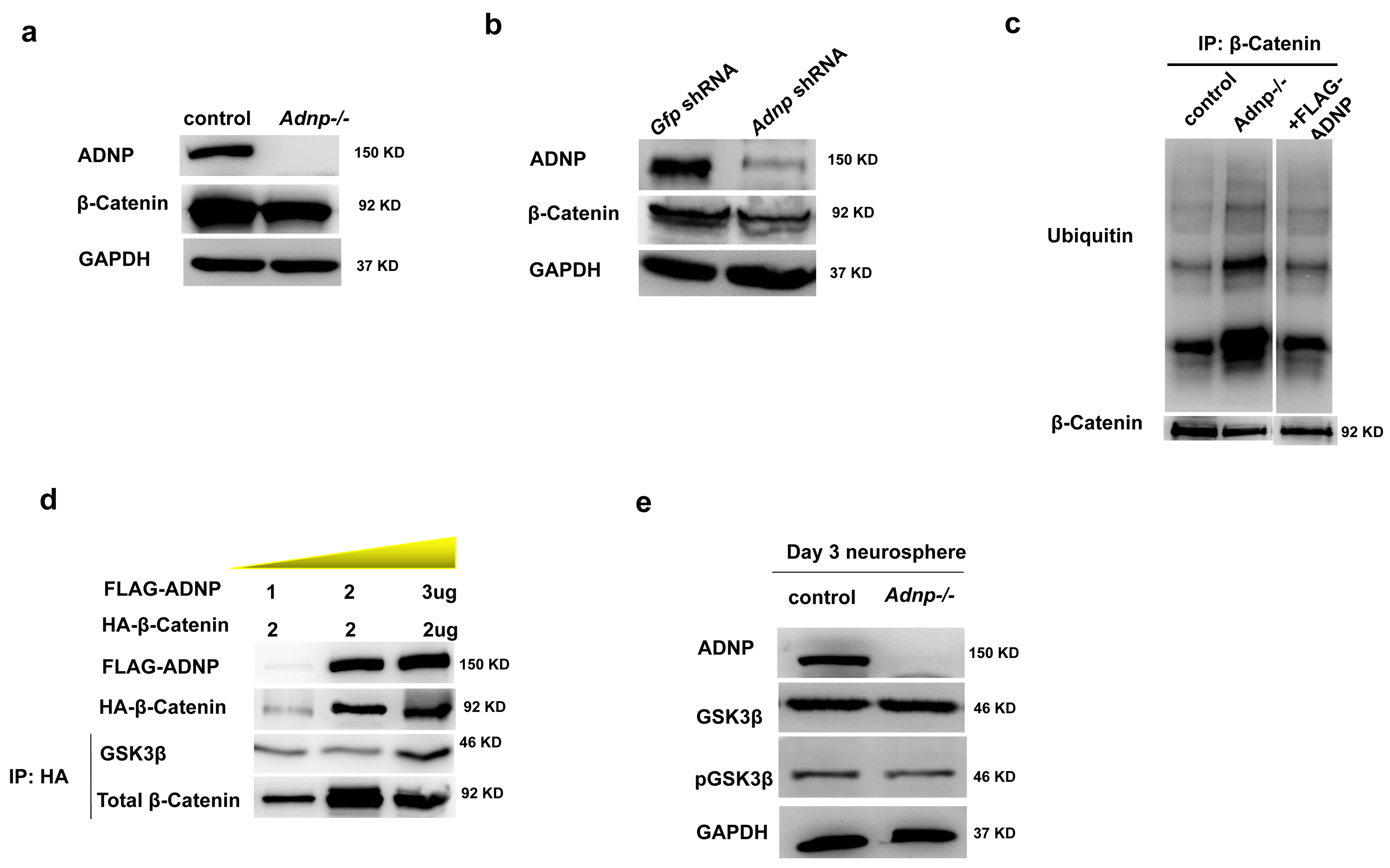
Related to Figure 6. **a** Representative WB showing the total β-Catenin levels in control and *Adnp-/-* ESCs. **b** WB showing the total β-Catenin levels in day 3 control and *Adnp* shRNA knockdown ESC-derived neurospheres. **c** Representative WB showing the ubiquitylation levels of β-Catenin in day 3 control, *Adnp-/-* and FLAG-ADNP overexpressing ESC-derived neurospheres. **d** Representative WB showing the effects of an increasing dose of ADNP on the co-transfected HA-β-Catenin and total β-Catenin levels. **e** Representative WB showing the effect of ADNP depletion on GSK3β and phosphorylated GSK3β levels. All WB experiments were repeated for two times.

**Supplementary Fig. 7.**
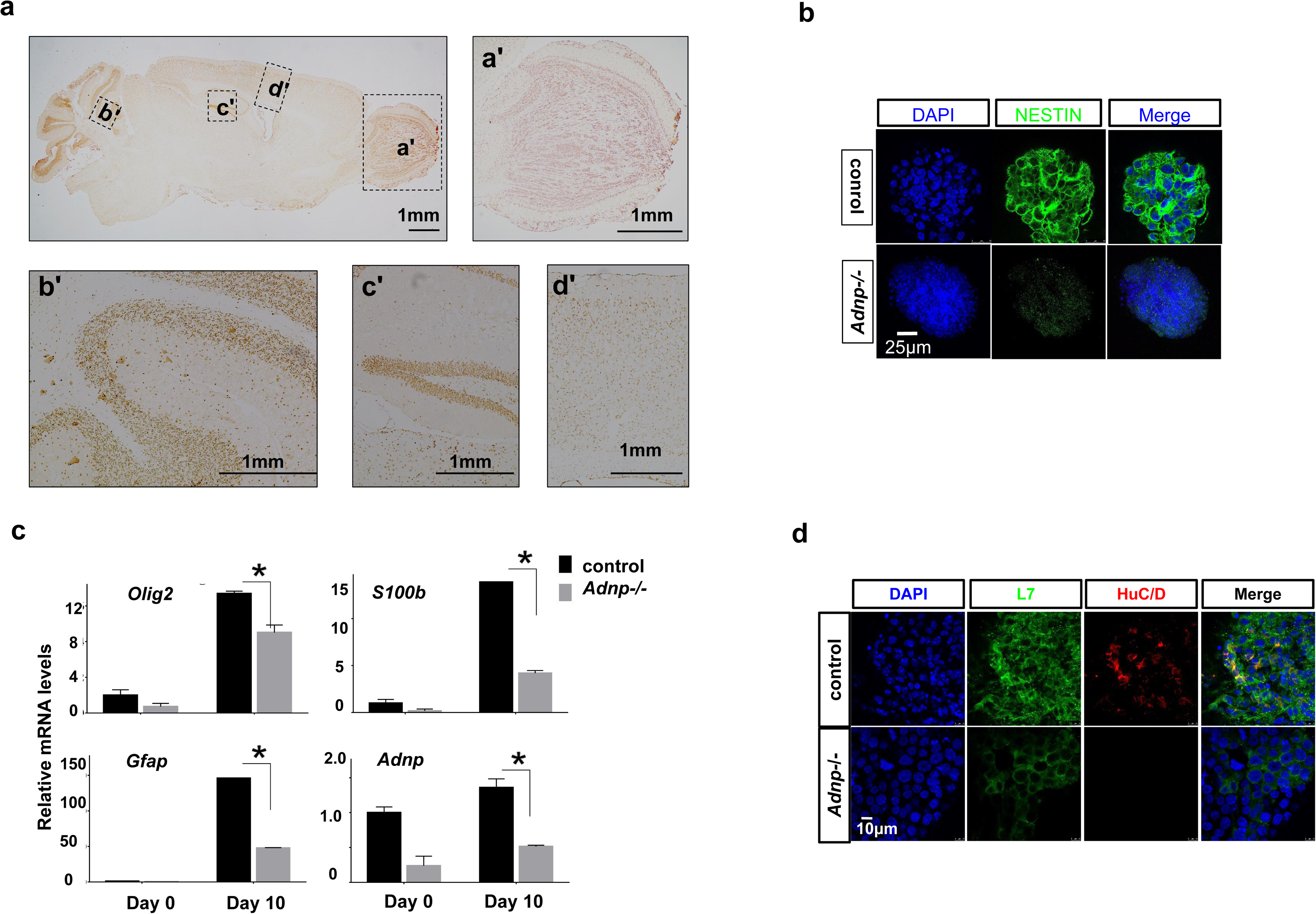
Related to Figure 8. **a** IHC image showing the expression of ADNP in 10-week old mouse brain. a, olfactory bulb, b, cerebellum, c, hippocampus, d, cortex. **b** IF staining of NESTIN for day 6 control and *Adnp*-/- ESC-derived SFEB aggregates. **c** The expression of the indicated neural genes in day 10 control and *Adnp*-/- ESC-derived EGL neural progenitors. **d** IF double-staining of L7 and HuC/D for day 18 control and *Adnp*-/- ESC-derived cerebellar cell types.

**Supplementary Fig. 8.**
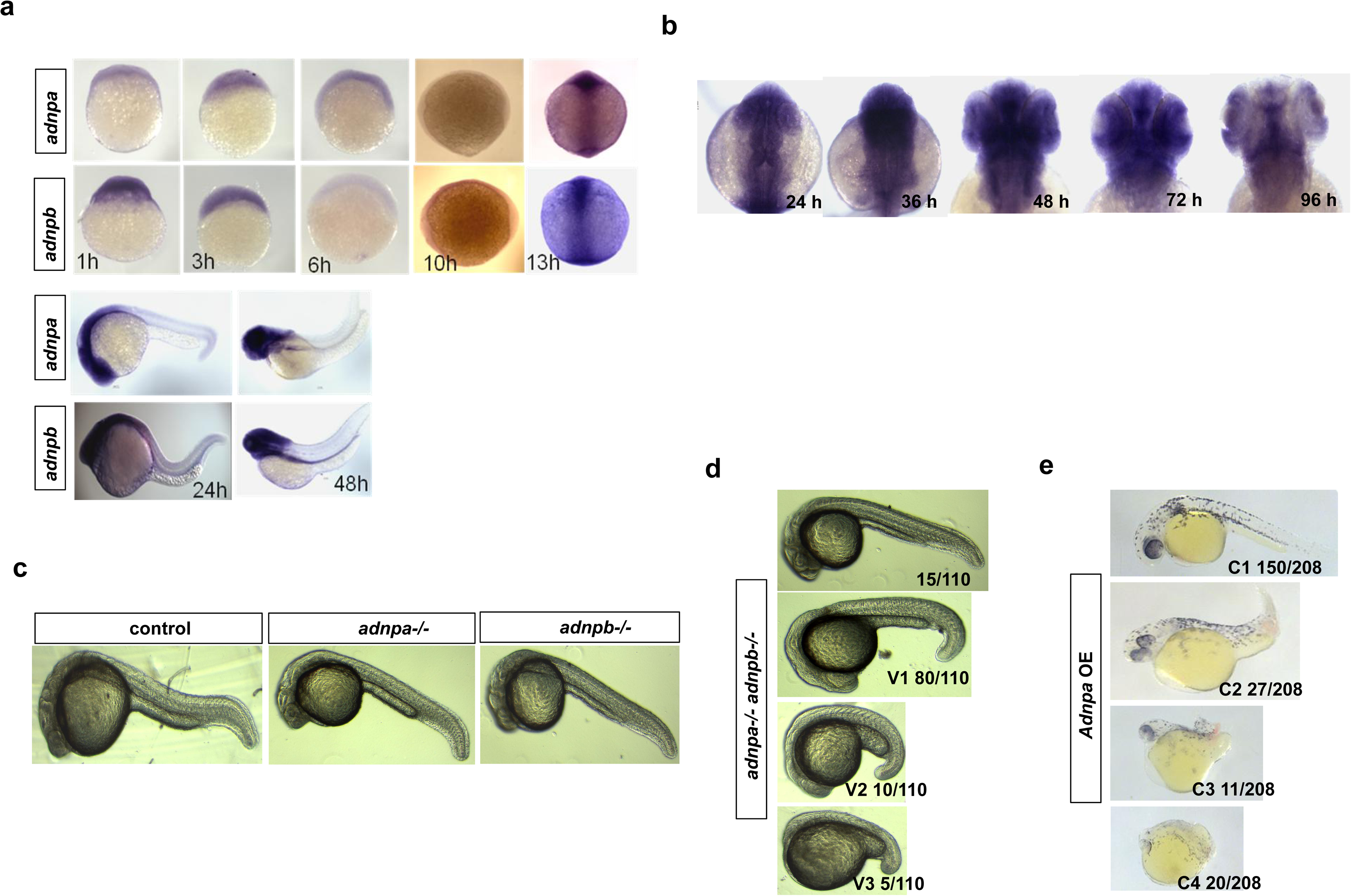
Related to Figure 10. **a** In situ hybridization showing the expression of *adnpa* and *adnpb* in embryos at different developmental stages. **b** In situ hybridization showing *adnpa* expression in head region of embryos at different developmental stages. Dorsal view of head region. **c** Representative morphology of 1 dpf control and *adnpa-/- adnpb-/-* zebrafish embryos. **d** Representative morphology of 1 dpf *adnpa-/- adnpb-/-* embryos. The ventralized phenotypes (V1-V3) were according to the DV patterning index^33^. **e** Representative morphology of 1 dpf *adnpa* overexpressing embryos. The dorsalized phenotypes (C1-C4) were according to the DV patterning index^33^

